# Regarding the *F*-word: the effects of data *Filtering* on inferred genotype-environment associations

**DOI:** 10.1101/2020.09.08.288308

**Authors:** Collin W Ahrens, Rebecca Jordan, Jason Bragg, Peter A Harrison, Tara Hopley, Helen Bothwell, Kevin Murray, Dorothy A Steane, John W Whale, Margaret Byrne, Rose Andrew, Paul D. Rymer

## Abstract

Genotype-environment association (GEA) methods have become part of the standard landscape genomics toolkit, yet, we know little about how to filter genotype-by-sequencing data to provide robust inferences for environmental adaptation. In many cases, default filtering thresholds for minor allele frequency and missing data are applied regardless of sample size, having unknown impacts on the results. These effects could be amplified in downstream predictions, including management strategies. Here, we investigate the effects of filtering on GEA results and the potential implications for adaptation to environment. Using empirical and simulated datasets derived from two widespread tree species to assess the effects of filtering on GEA outputs. Critically, we find that the level of filtering of missing data and minor allele frequency affect the identification of true positives. Even slight adjustments to these thresholds can change the rate of true positive detection. Using conservative thresholds for missing data and minor allele frequency substantially reduces the size of the dataset, lessening the power to detect adaptive variants (i.e. simulated true positives) with strong and weak strength of selections. Regardless, strength of selection was a good predictor for GEA detection, but even SNPs under strong selection went undetected. We further show that filtering can significantly impact the predictions of adaptive capacity of species in downstream analyses. We make several recommendations regarding filtering for GEA methods. Ultimately, there is no filtering panacea, but some choices are better than others, depending largely on the study system, availability of genomic resources, and desired objectives of the study.

## Introduction

Identifying genomic patterns associated with adaptation in wild populations can provide information to support management strategies as well as facilitate fundamental discoveries (Garner et al., 2016; Sgrò, Lowe, & Hoffmann, 2011). We can improve our understanding of the response of species to changing climates and their evolutionary potential by leveraging knowledge about adaptive genetic variation in natural populations (Browne, Wright, Fitz-Gibbon, Gugger, & Sork, 2019; Razgour et al., 2019; Sork, 2017). Genotype–environment association (GEA) methods are used to identify potentially adaptive loci in non-model systems based on correlations between allele frequencies and environmental data. In recent years, there has been a proliferation of genomic studies on landscape adaptation using GEA analyses (Ahrens et al., 2018), which is becoming a standard part of the analytical pipelines for landscape genomics.

The utility of GEA analyses is limited by several problems, including the presence of false positives (type I errors) (Storz, 2005). While, false negatives (type II errors) are likely common due to controlling for population structure (Sork et al., 2013), they are unlikely to limit or confound the GEA results. False positives are present in GEA outputs regardless of filtering, significance thresholds or false discovery corrections (Forester et al., 2018). From a biological perspective, false positives are genomic variants significantly associated with the environment through random, neutral processes. For example, demographic processes can generate clines in allele frequencies that covary with environmental gradients, leading to neutral SNPs potentially being falsely identified as adaptive (François, Martins, Caye, & Schoville, 2016; Hoban et al., 2016; Lotterhos & Whitlock, 2015). However, these impacts will vary depending on the unique demographic history (e.g. bottlenecks, population growth, or rapid expansion) of a species. Many GEA methods control for patterns of population structure, to reduce false positive call rates, but by doing so, true positives are also at risk of becoming false negatives (Nadeau, Meirmans, Aitken, Ritland, & Isabel, 2016; Orsini, Mergeay, Vanoverbeke, & Meester, 2013). One way to control for false positive call rates is to combine the results of multiple approaches in the hope of identifying loci with well-supported associations with environmental variables (Meirmans, 2015). However, the outcomes of these approaches are variable (Nadeau et al., 2016) and, this is not surprising given the numerous statistical models and methods used to mitigate the confounding effects of genetic structure. Also, the consequences of false positives could vary, depending on the conservation or management applications associated with the analysis. For example, the presence and overrepresentation of false positives could have implications for conservation actions, through the identification of patterns of putative adaptation that are supported more by false positives than true positives (i.e. the noise is stronger than the signal).

The occurrence of false positives is partially attributable to incomplete genome sampling (Lowry et al., 2017). The proportion of the genome sampled can be influenced at many stages of the workflow, including choice of genotyping method, library preparation method (e.g. enzyme choice), bioinformatic processing, and data quality filtering (O’Leary, Puritz, Willis, Hollenbeck, & Portnoy, 2018). Most GEA studies of non-model organisms employ reduced representation approaches, as they are cost-effective, do not require extensive genomic resources (e.g. reference genomes) (Manel et al., 2016) and often yield thousands of loci scattered across a species’ genome. Yet, even small genomes are poorly sampled through reduced representation library preparation. For example, a dataset of 20 k SNPs only represents ∼0.7% of a 550 Mbp genome with a linkage disequilibrium decay of 200 bp (2.75 million linkage blocks). Thus, for many reduced representation approaches, the likelihood of detecting positive associations is limited by querying a very small proportion of the genome. Previous studies have amply reviewed how choices made during library preparation and bioinformatic processing impact the level of genome sampling that can be achieved for any given reduced representation dataset (Mastretta-Yanes et al., 2015; O’Leary et al., 2018). In addition, total sample size is also known to have an impact on the power of GEA analyses and identification of false positives (Lotterhos & Whitlock, 2015). While the importance of sample size alone has been discussed previously as an important factor for sample design for GEA analyses (Lotterhos & Whitlock, 2015; de Mita et al., 2013), it is unknown how sample size interacts with filtering choices. Therefore, in this study we explore the explicit impact of data quality filtering on downstream GEA results.

Filtering remains incredibly challenging, and a highly important aspect of population genomics data analysis (Andrews & Luikart, 2014). Optimal, default filtering settings suitable for all GEA studies are unlikely, given the range of organisms and research questions explored. Even so, documenting the effects of data filtering on analyses has proved highly useful for other population genetic applications, assisting researchers to set filters that are appropriate for their experimental design and individual study goals (Narum, Buerkle, Davey, Miller, & Hohenlohe, 2013). For example, it has been shown previously that SNP calling and filtering settings can affect estimates of heterozygosity and *F*_ST_ (Díaz-Arce & Rodríguez-Ezpeleta, 2019; Shafer et al., 2017), routinely used in conservation decision making (Gautier et al., 2012; Pool, Hellmann, Jensen, & Nielsen, 2010). Minor allele frequency (MAF) filtering settings can change *F*_ST_ estimates (Hendricks et al., 2018; Linck & Battey, 2019), due to the inclusion of locally isolated alleles increasing the perceived dissimilarity of populations. Liberal thresholds of missing data have been shown to reduce estimates of expected heterozygosity and increased inference of inbreeding; however, the results vary across species (Fu, 2014). Stringent filtering increases completeness of the dataset at the expense of the number of SNPs retained and the proportion of the genome sampled. While it is general practice to filter missing data to low levels, no studies to date, as far as we are aware, have investigated the impact of missing data on downstream GEA results. In addition, filtering of reduced representation datasets from organisms without genomic resources is even more critical, because *de novo* alignment can introduce errors (O’Leary et al., 2018). While the importance of filtering has been acknowledged, the impacts of filtering thresholds on GEA analyses have yet to be fully investigated.

In many cases, GEA analyses and outputs are cited as being useful for downstream applications, including the improvement of management, conservation, and breeding programs. While commendable, we do not know how filtering choices might impact final recommendations. As the dataset changes due to filtering, so too will the identified set of putatively adaptive SNPs, and these differences could be compounded when extrapolating across environmental space. Often these extrapolated maps, of adaptive genomic variation across species’ ranges, are the currency of interpretation for stakeholders and decision-makers. The connections between geospatial predictions of adaptation and genomic variation to support management / conservation outcomes is evident in studies on birds (Bay et al., 2018) and grasses (Ahrens et al., 2020), where researchers quantify the heterogeneity of genomic vulnerability to climate change. However, these predictive outputs could be affected by filtering choices.

Filtering requires subjective decisions about how best to compile the best available dataset to investigate genomic adaptation across landscapes, while limiting the proportions of false positives and false negatives identified by GEA analyses. No definitive filtering guidelines for GEA currently exist. Instead researchers are left to iteratively change filtering thresholds and subjectively choose a perceived optimal dataset for the question at hand (as demonstrated by the range of filtering settings identified in a GEA meta-analysis; Ahrens et al., 2018). This subjective process may result in ambiguous interpretation and the potential for bias in the reporting of results. As the incorporation of GEAs into analytical pipelines increases, it is important to establish objective guidelines to assist researchers in determining the impact that filtering can be expected to have on downstream GEA results. Therefore, we ask two questions: 1) how does filtering affect the identification of putatively adaptive loci? and, 2) how does our ability or inability to identify associations affect downstream applications? To answer these questions, we test four common assumptions:

1. More stringent filtering reduces identification of false positives.
2. Loci with strong selection strengths will be identified as significant, regardless of filtering choices.
3. Combining GEA analyses reduces false positive call rates.
4. Extrapolation of adaptive variants across the landscape reveals consistent areas of climate adaptation.

We test these assumptions using both empirical and simulated data sets, the latter matched to the empirical demographic scenarios with the addition of known true positives. We explore how early filtering decisions affect conservation and management decisions and provide guidelines for data filtering to optimise the effectiveness of GEA methods.

## Methods

### SNP and climate data

We chose two reduced representation SNP datasets from different genera within the eucalypt group: *Eucalyptus microcarpa* (Maiden) Maiden (Jordan, Hoffmann, Dillon, & Prober, 2017) and *Corymbia calophylla* (Lindl.) K.D.Hill & L.A.S.Johnson (Ahrens, Byrne, & Rymer, 2019). Both species are native to south-eastern and south-western Australia respectively (Figure 1). By comparing phylogenetically close species, we minimised potential confounding effects arising from using species with very different genomes, thereby allowing us to focus on how filtering affects GEA results.

**Figure 1.**
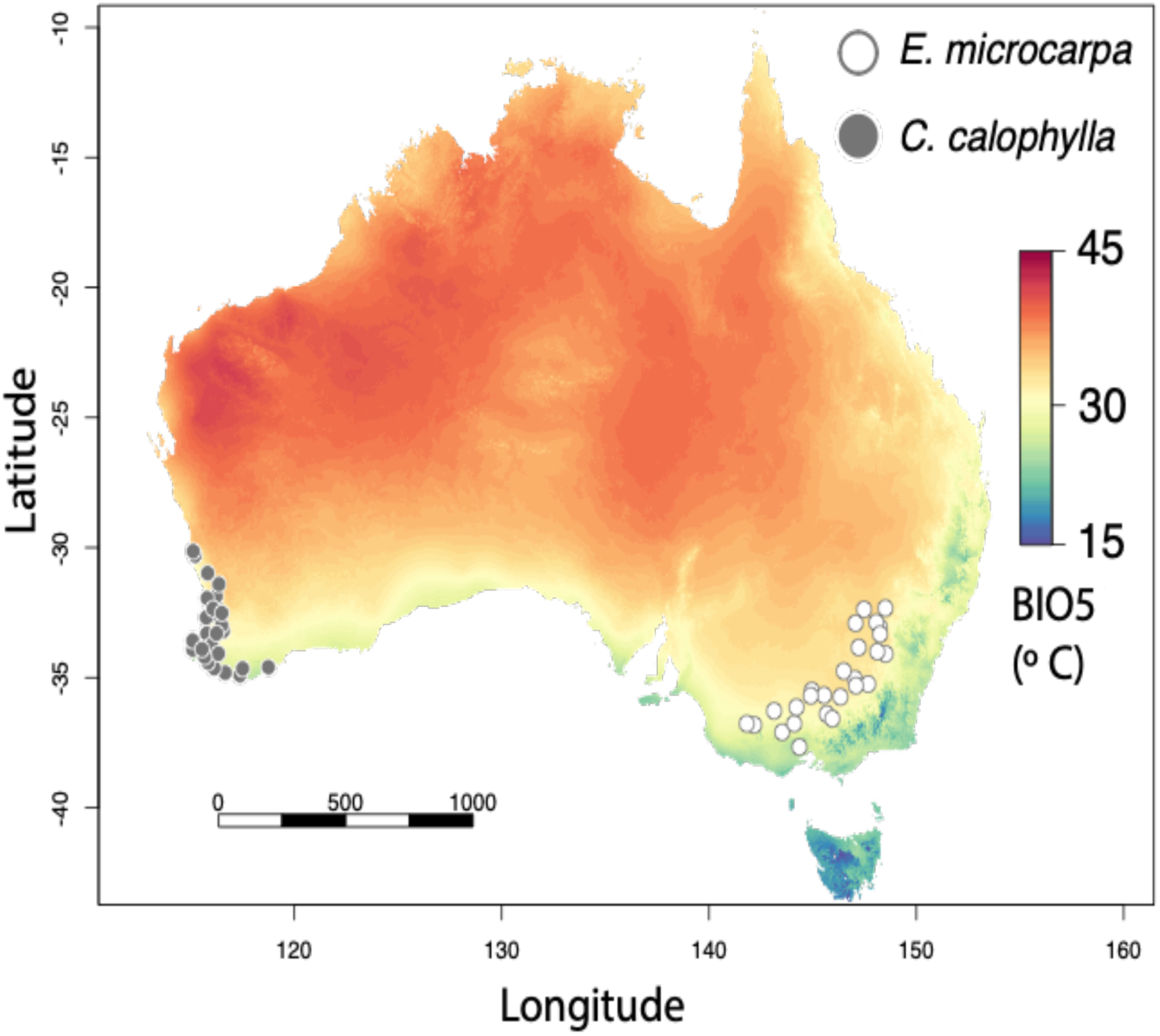
Map of the sampled locations for the two study species with maximum temperature of the warmest month (BIO5) shown across Australia.

The datasets were based on sampling across the range of each species. The *E. microcarpa* dataset consisted of a total of 577 samples from 26 populations and the *C. calophylla* dataset comprised 263 samples from 27 populations. Genomic data for both species were generated using DArTseq (Diversity Arrays Technology P/L, Canberra, Australia), with the same library preparation, multiplexing, and sequencing protocols. The raw, unfiltered genotype data were used as the input datasets, with different filtering applied as described below. Genotypes were quality filtered prior to analysis, retaining those with an individual minimum read-depth of 10x, minimum genotype quality Phred-score of 30 and a maximum mean read-depth of 100x, retaining only biallelic SNPs.

Climate data were extracted from WorldClim (Fick & Hijmans, 2017) for each sampling location using the R package *raster* (R core team 2019). We chose the mean maximum temperature of the warmest month (BIO5) to test the effect of filtering on genotype-environment association (GEA) analyses. Temperature was selected as it is commonly used in GEA analyses and a key selective force given projected increases into the future; BIO5 represents the high temperature extremes, presumably a greater selective pressure than mean annual temperatures in Australia (Prober et al., 2016; Costa e Silva, Potts, Harrison, & Bailey, 2019). To assess the potential effect of multiple variables confounding GEA results, we also tested mean precipitation of the driest month (BIO14), representing a second key selective force of precipitation. Assessments of spatial autocorrelation (Moran’s *I*) and effective population size, given the environment (*n*_eff-env_) was performed, provide critical metrics for determining which climate variables have greater power to detect SNPs under selection (details in Supplementary information).

### Simulated data set creation

Simulated SNP datasets were generated to be comparable to the empirical datasets, with two main motivations. First, the effect of missing data can be studied by generating complete simulated SNP datasets, and then implementing different levels of ‘missingness’. Second, simulated datasets enable evaluation of the performance of GEA (rates of detection of false positives and negatives) in relation to different filtering treatments. This can be accomplished by including known true positives (TP) with different magnitudes of selection pressure.

Simulated datasets were generated using the *simulate*.*baypass* R function in BayPass (Gautier, 2015). This function creates simulated datasets under a BayPass model (see Coop, Witonsky, Rienzo, & Pritchard, 2010; Günther & Coop, 2013) using an empirical matrix of allelic covariances (the Ω matrix). It generates SNPs whose allele frequencies vary across populations according to the covariance matrix previously estimated from the empirical datasets, with an additional associations of prescribed strength to a bioclimatic variable. Two simulated datasets were generated based on the species’ empirical data, hereafter referred to as ‘*Sim microcarpa’* and ‘*Sim calophylla*’ to distinguish from empirical datasets of *E. microcarpa* and *C. calophylla*, respectively.

We simulated population-level allele counts for ∼25 000 ‘neutral’ SNPs plus 200 ‘adaptive’ (i.e. simulated SNPs that are correlated with a specific climate variable) SNPs whose coefficients of association with each of the two bioclimatic variables were drawn from a uniform distribution between −0.3 and 0.3 (beta.coef). We chose these selection coefficients knowing that, at their extremes, they are likely greater than the values we would find in wild populations. We did this intentionally to verify that loci with very strong selection coefficients were highly likely to be identified in the GEA analyses. Other *simulate*.*baypass* parameters were chosen so that the simulated data resembled our empirical datasets. For example, the simulation function uses a beta distribution to describe the frequencies of ancestral alleles among loci. We chose the parameters for this distribution by fitting the beta distribution to the minor allele frequencies observed in the empirical datasets. *Corymbia calophylla* returned shape1 = 0.54 and shape2 = 0.53, whereas *E. microcarpa* returned shape1 = 0.43 and shape2 = 0.43. Fixed loci were removed from the simulated datasets, resulting in a loss of 1000-1600 SNPs per dataset. We also wanted to approximate, in the simulations, the way missing data were distributed across samples and across loci in the empirical data sets. We therefore began by fitting statistical distributions to frequencies of missing genotypes across loci and samples in the empirical data. We used the estimated distributions to impose missing alleles on the loci and samples across the simulated datasets (Figure S1). If we sampled from a distribution and obtained a negative number of missing genotypes for a locus, we set the value of missingness for that locus to 0.

### Subsetting datasets

To understand how filtering choices affect the ability of GEAs to identify true positives, we filtered each data set by minor allele frequency (MAF), missing data (MD), and the number of samples per population (all 150 data sets represented in Table 2). We chose three MAF to explore (0.01, 0.05, and 0.1; Table 2) based on the most commonly applied thresholds (Ahrens et al., 2018). We applied five MD thresholds (10%, 20%, 30%, 40%, and 50%; Table 2). The most commonly applied MD thresholds are between 10 and 30%; we included thresholds up to 50% to test how less-stringent MD thresholds would behave with GEA methods.

**Table 1.** Attributes of empirical and simulated datasets. Pearson’s correlation coefficient (*r*); spatial autocorrelation (Moran’s *I*); effective sample size due to environment (*n*_eff-env_); BIO5 – maximum temperature of the warmest month; BIO14 – precipitation of the driest month; number of SNPs remaining after filtering for largest and smallest analysis datasets (#SNP).

| | Empirical | | | | Simulation | | | Structure | $r$ | Moran's $I$ | $n_{\text{eff-env}}$ |
| --- | --- | --- | --- | --- | --- | --- | --- | --- | --- | --- | --- |
| species | # samples | # pops | #SNP largest | #SNP smallest | # samples | #SNP largest | #SNP smallest | $F_{\text{ST}}$ | BIO5 ~ BIO14 | BIO5 | BIO5 |
| *E. microcarpa* | 577 | 26 | 25,826 | 2,931 | 650 | 20,685 | 3,494 | 0.01 | 0.014 | 0.267 | 15.1 |
| *C. calophylla* | 263 | 27 | 25,811 | 5,595 | 270 | 21,255 | 5,031 | 0.05 | -0.75 | 0.327 | 13.6 |

**Table 2.**
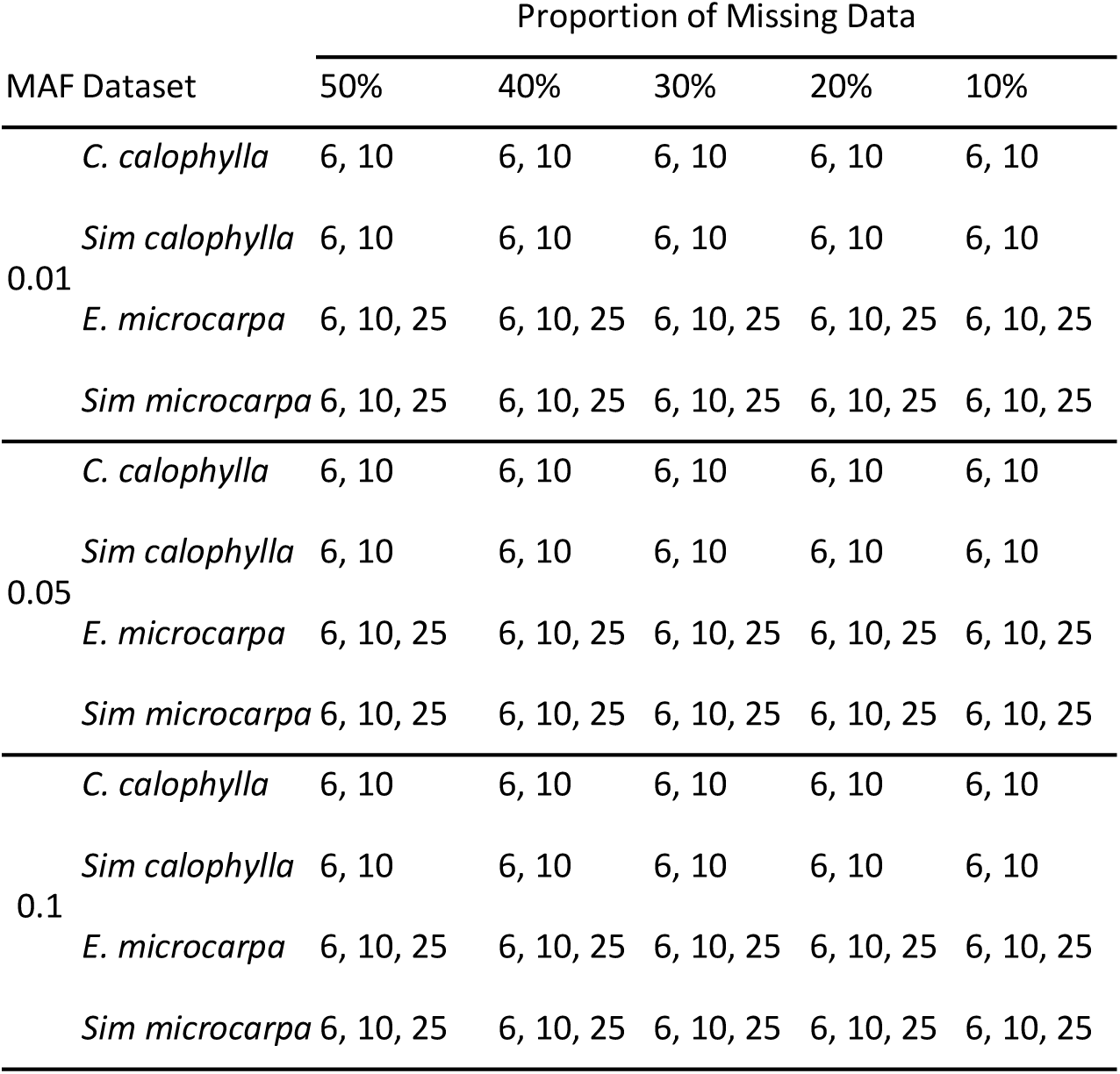
Matrix detailing the 150 data filtering combinations explored in the present study. Numbers within the table represent the total number of individuals per dataset: 6, 10, or 25 individuals per population. The number of populations remained constant throughout the study (*C. calophylla* – 27 populations; *E. microcarpa* – 26 populations). MAF – minor allele frequency.

The importance of biological sampling design on GEA analyses has been demonstrated previously (see Forester et al., 2018; Lotterhos & Whitlock, 2015; de Mita et al., 2013 for more thorough treatments of sampling design), and do not try to replicate these studies but rather. Six individuals per population is often regarded as the minimum sample size for population genetics analyses when thousands of SNPs are available (Nazareno, Bemmels, Dick, & Lohmann, 2017; Willing, Dreyer, & Oosterhout, 2012) and GEA studies (Lotterhos & Whitlock, 2015). We therefore tested the effect of using 6 or 10 individuals per population for both species, as well as 25 individuals per population for *E. microcarpa*, reflecting the empirical *C. calophylla* and *E. microcarpus* datasets, respectively.

### GEA Analyses

We focused on three commonly used GEA methods with different underlying computational models to identify SNP-climate associations. We compared two univariate methods, LFMM2 (Caye, Jumentier, Lepeule, & François, 2019) and BayPass (Gautier, 2015) which associates each SNP individually with a given climate variable, and one multivariate method, redundancy analysis, RDA following the usage in Forester et al., (2018).

LFMM2 uses discrete ancestral clusters computed via principal component analysis (PCA) to control for population structure, and a least-squares approach for confounder estimation of genomic data (Caye et al., 2019). As LFMM2 requires a full data set, we imputed data using the mean method with the *impute* function as the default and may be considered the ‘worst case scenario’ imputation method (note: the mean method is a naive imputation method, and we suggest using other imputation methods). We ran PCAs for each data set to assess the change in population structure as a result of filtering choices (data not shown). As expected, population structure varied across datasets, likely due to the low, but present population structure (*F*_ST_ = 0.05 & 0.01; Table 1). We observed only very slight changes from a *K* = 3 to a *K* = 4, 5, or 6, with *K* = 3 being the most consistent solution for both species. Therefore, we used *K* = 3 for all LFMM2 analyses to allow direct comparisons across data sets. Significant associations were called at α = 0.001 after applying a false discovery rate as suggested by Caye et al., (2019). We explored lower significance thresholds but found they were too permissive, returning high numbers of false positives; 0.001 seemed to be similar to the BayPass significance factor, a Bayes Factor (BF), of 20.

BayPass uses an Ω matrix to account for population structure based on allelic covariance between populations. BayPass analyses were run following the methods described in the BayPass manual. We ran the standard model twice to obtain the Ω matrix, and averaged the Ω matrix across runs. The mean Ω matrix was used as the covariance matrix within the auxiliary model, which calculates a BF to assist with identification of SNP-climate associations. The auxiliary model was run twice, and results averaged across runs. The parameters used for both models (standard and auxiliary) were 20 pilot runs for 1000 iterations, 2500 burn-in, and 1000 MCMC samples. Significant associations were called at a BF > 20, considered ‘decisive’ evidence (Jeffreys, 1961). As above, for LFMM2, we explored other significance levels with results returning high numbers of false positives.

Complementing the univariate GEA analyses, we also performed a redundancy analysis (RDA). This multivariate method has been shown to be robust across a wide range of selection strengths, demographic histories, sampling designs, and in the presence of many levels of population structure (Forester et al., 2018).To address the RDA requirement of a complete data set, we calculated and used population-level allele frequencies, instead of imputation. For RDA, an α = 0.05 was used to extract significant SNPs along the two climate axes, across the three main RDA axes. Variance inflation factors (VIF) were used to check multicollinearity between the two climate variables, *C. calophylla* returned 2.35 VIF for both climatic variables and *E. microcarpa* returned 1.00 VIF for both, indicating that these are sufficiently independent to identify associations via RDA because they are below 10 (Zuur et al., 2010).

For each dataset and analysis, we recorded the SNPs that were identified as having significant associations with environment. For simulated datasets, we recorded which SNPs were true positives (TP) and which were false positives (FP). We also recorded ‘pseudo positives’ (PP), defined as SNPs that were found to be significantly associated with one climate variable but were in fact TP for the other climate variable i.e. were identified as significantly associated with BIO5 but were actually adapted to BIO14.

In order to test whether there is a strength of selection threshold for which GEA methods achieve a 100% TP call rate, we plotted strength of selection (beta coefficient applied during simulations) against the significance of association for BayPass (BF) and LFMM2 (calibrated *P*-value) for *Sim calophylla*. We also calculated the difference between the significance values for each MAF threshold and the standard deviation. This estimate allowed us to quantify the mean differences and variance between data sets differentiated only by MAF.

### Impacts of filtering on extrapolation and interpretation of adaptive variation

To determine how filtering thresholds may affect the downstream extrapolation of putatively adaptive genomic variation across geographic space, we estimated the genomic-informed ‘climate selection surface’ for both species. Here, a climate selection surface refers to the prediction of adaptation through geographic space. This extrapolation followed the logic of Steane et al. (2014), but using RDA instead of canonical analysis of principal coordinates (details provided in supplementary information). The effect of each filtering parameter was explored separately in the simulated datasets, holding other filtering parameters constant (e.g., when assessing the effect of MAF, the MD and sample size thresholds were held constant). We also compared the impact of different filtering methods on the empirical datasets for the most liberal (MD = 50%; MAF = 0.01) and conservative (MD = 10%; MAF = 0.1) datasets. Significant differences between climate selection surfaces were determined using a pixel pairwise z-score test. Here, the liberal dataset was compared to the conservative dataset, such that a positive difference between the two resulting climate selection surfaces corresponds to the liberal dataset predicting more adaptive variation, and a negative difference corresponds to the conservative dataset predicting more adaptive variation.

## Results

### Effects of filtering on GEA outputs – simulated data

Using simulated data that reflected natural population structure and climate gradients across *C. callophylla* and *E. microcarpa* (*‘Sim calophylla’* and *‘Sim microcarpa’* respectively), we found that data filtering influenced the identification of ‘adaptive’ SNPs. Filtering regimes differentially impacted the data sets and GEA programs in various ways. Both filtering thresholds (missing data (MD), minor allele frequency (MAF)) and biological sample size influenced the number of significant SNP-climate associations. Furthermore, filtering thresholds also impacted the number of true positives (TP), false positives (FP) and pseudo-positives (PP).

With the exception of RDA for *Sim calophylla*, the GEA methods identified SNP associations with BIO5, including TPs (Figures 2 & 3). The multivariate RDA approach performed exceedingly poorly for *Sim calophylla* and only moderately well for *Sim microcarpa* compared to the other two GEA methods in all aspects, particularly in identifying TPs. For *Sim calophylla*, this finding was surprising and might be due to the fact that the climate variable is closely associated with the population structure (see Ahrens et al., 2019 for details), identifying all TPs as false negatives; alternatively, the demographic history *C. calophylla* may make RDA less sensitive to true associations, as no associations were found in the empirical dataset either. Because of this complication, we focus the results on BayPass and LFMM2.

**Figure 2.**
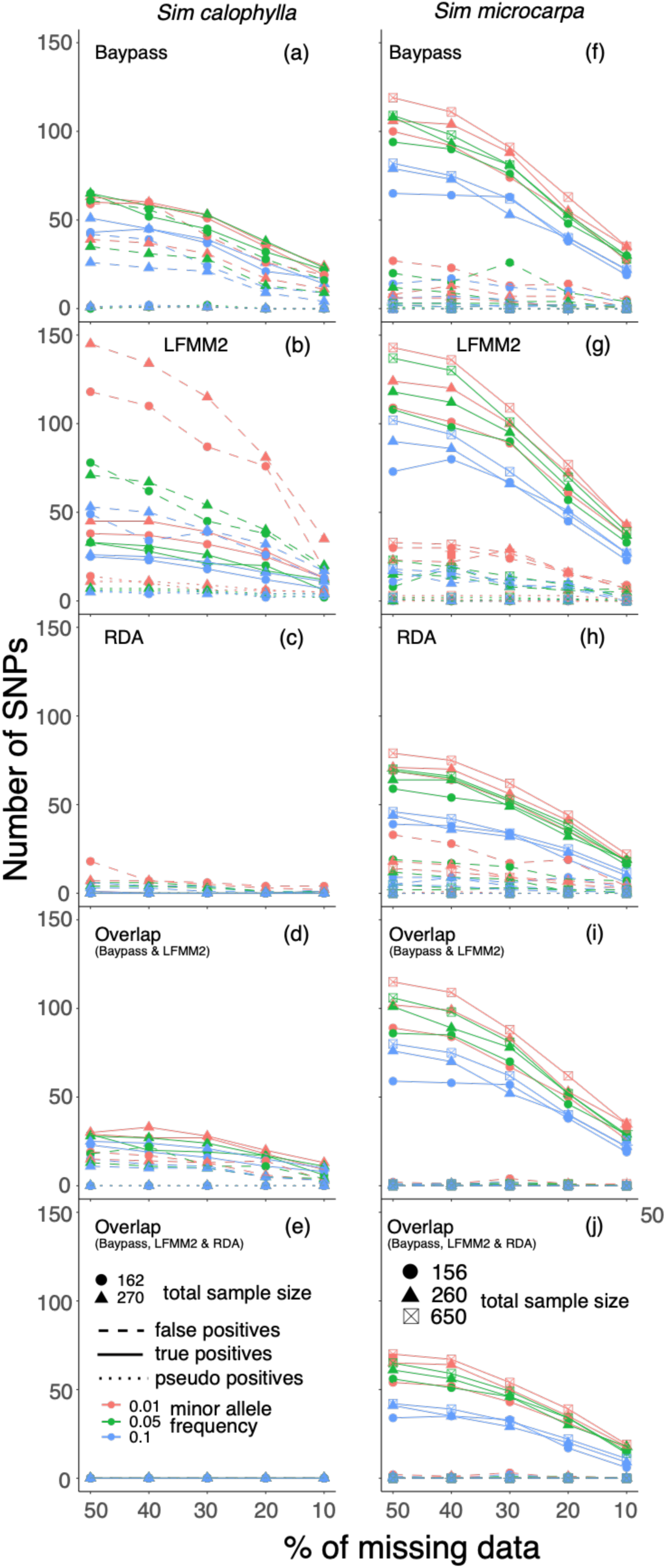
The number of significant SNP-climate associations with BIO5 (maximum temperature of the warmest month), for the simulated datasets (a-e) *Sim calophylla* and (f-j) *Sim microcarpa* using three GEA analytical approaches: (a,e) BayPass, (b,g) LFMM2, and (c,h) RDA; including ‘overlap’ – common identified associations – between (d,i) BayPass and LFMM2, and (e,j) BayPass, LFMM2 and RDA. Associations called false positives (FP) – significant ‘non-adaptive’ SNPs; true positives (TP) – significant ‘adaptive’ SNPs; and pseudo positives (PP) – SNPs ‘adaptive’ for BIO14 (precipitation of the warmest month) but found to be significantly associated with BIO5.

The numbers of TPs and FPs increased with higher proportions of missing data (Figure 2). This pattern reflects, in part, the total number of SNPs retained in each filtered dataset, with fewer SNP-climate associations and TPs retained when more stringent filtering was applied (Figure S2). There were significant relationships between the number of TPs found and the total number of SNPs kept in the analysis for both species (*Sim microcarpa* – *r*^2^ = 0.93, *p* = <0.0001; *Sim calophylla* – *r*^2^ = 0.89, *p* = <0.0001) (Figure S2). On the other hand, the amount of missing data had little influence on the proportion of TPs in ‘All Associations’ (AA) and, thus, the ratio of TPs to AAs remained constant within method and species (TP:AA; Figure 3). Although the TP:AA ratio was markedly different between species and between methods within species (Figure 3).

**Figure 3.**
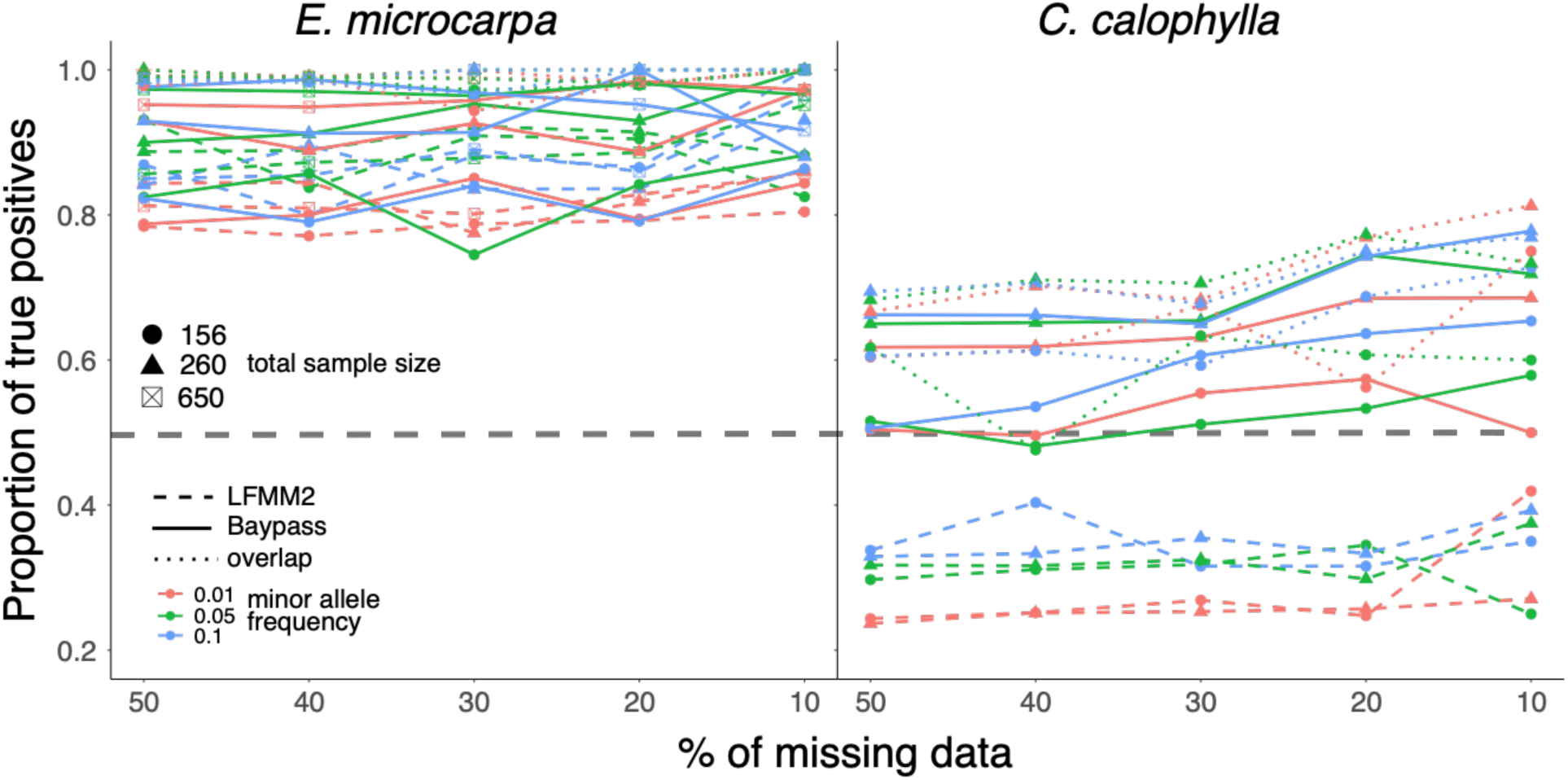
The proportion of *True Positives* (TP) among all identified associations (AA) called in BayPass, LFMM2, and the SNPs shared between them. The dashed horizontal line indicates 50% TPs in AA; equal to a 1:1 ratio of TPs vs false positives (FP). For values above this line TPs > FPs, while below the line TPs < FPs.

In general, a smaller MAF identified more TPs and more FPs than a large MAF (Figure 2). The increase in FPs was especially apparent in the LFMM2 analysis for the *Sim calophylla* data, where a MAF of 0.01 yielded nearly twice as many FPs as TPs (Figure 2b). For the *Sim microcarpa* dataset, a MAF of 0.1 identified substantially fewer TPs then lower MAFs, although the decrease in FPs was not as clear because of the already low FP call rate. The proportion of TPs in AAs varied with MAF (Figure 3). For the *Sim calophylla* data, a larger MAF generally resulted in a higher proportion of TPs (higher ratio of TP:AA). For the *Sim microcarpa* data, a MAF of 0.01 generally had the lowest proportion of TPs (lowest ratio of TP:AA), with the highest proportion of TPs varying between MAF 0.05 and 0.1 depending on the program used and amount of missing data.

Sample size and pseudo positives differed between species and method. Larger biological sample sizes consistently identified more TPs for *Sim microcarpa*, whereas sample size had less influence on TP identification (Figure 2; more detailed results about sample size are in the supplementary information). Pseudo positives (PP) were at or near zero for *Sim microcarpa* for both BayPass and LFMM2, but PPs were detected for *Sim calophylla* in LFMM2, but few in BayPass (Figures 2a).

### Overlapping results

A common approach for determining putatively ‘adaptive’ SNPs is to select those SNPs identified in multiple, independent analyses (Lotterhos et al., 2017), the rationale being that these SNPs are more likely to be TPs. Our results show a slight increase in the proportion of TPs identified (increased TP:AA) when results from independent analyses were combined (Figure 3). This was due to a small reduction in the number of FPs compared to the most conservative method (i.e. BayPass). However, this reduction in FPs came at the cost of fewer TPs being retained. In general, the number of TPs retained was reduced to the level of the more conservative dataset. For *Sim microcarpa*, the number of TPs was reduced to BayPass numbers for the BayPass-LFMM2 overlap (Figure 2i) and reduced to RDA numbers for the BayPass-LFMM2-RDA overlap in *Sim microcarpa* (Figure 2j). *Sim calophylla* had a substantially greater decrease in TPs when comparing the overlap between BayPass and LFMM2, dropping to less than either Baypass or LFMM2 (Figure 2d). There were no identified TPs common to all three analyses for *Sim calophylla*, reflecting the lack of TPs from RDA (Figure 2e). Using multiple methods decreased the number of FPs, to the point of there being very few or zero FPs for *Sim microcarpa* (Figure 2). This decrease in FPs compared to TPs when using multiple methods slightly increased the proportion of TPs in the set of SNPs common to multiple GEA methods (Figure 3).

### The influence of selection strength on identifying associations

We hypothesised that the strength of selection prescribed in the simulations would influence the magnitude of the association inferred, and ultimately, the likelihood of detecting TPs. In particular, we wanted to know if it was possible to identify a threshold above which the call rate for TP was 100%. The strength of selection for individual TPs did impact the identification of significant associations. While never 100% accurate at any strength of selection, linear models revealed significant relationships between the strength of selection and levels of significance for BayPass and LFMM2 (r^2^ = 0.47 and 0.22 respectively), showing the strength of selection does have some effect on results (Figure S3). However, a threshold for high TP call rates was only observed at low levels of missing data (10%) for BayPass (strength of selection +/− 0.28; Figure 4). This threshold disappeared when we included more SNPs through filtering and no threshold was identified for LFMM2.

**Figure 4.**
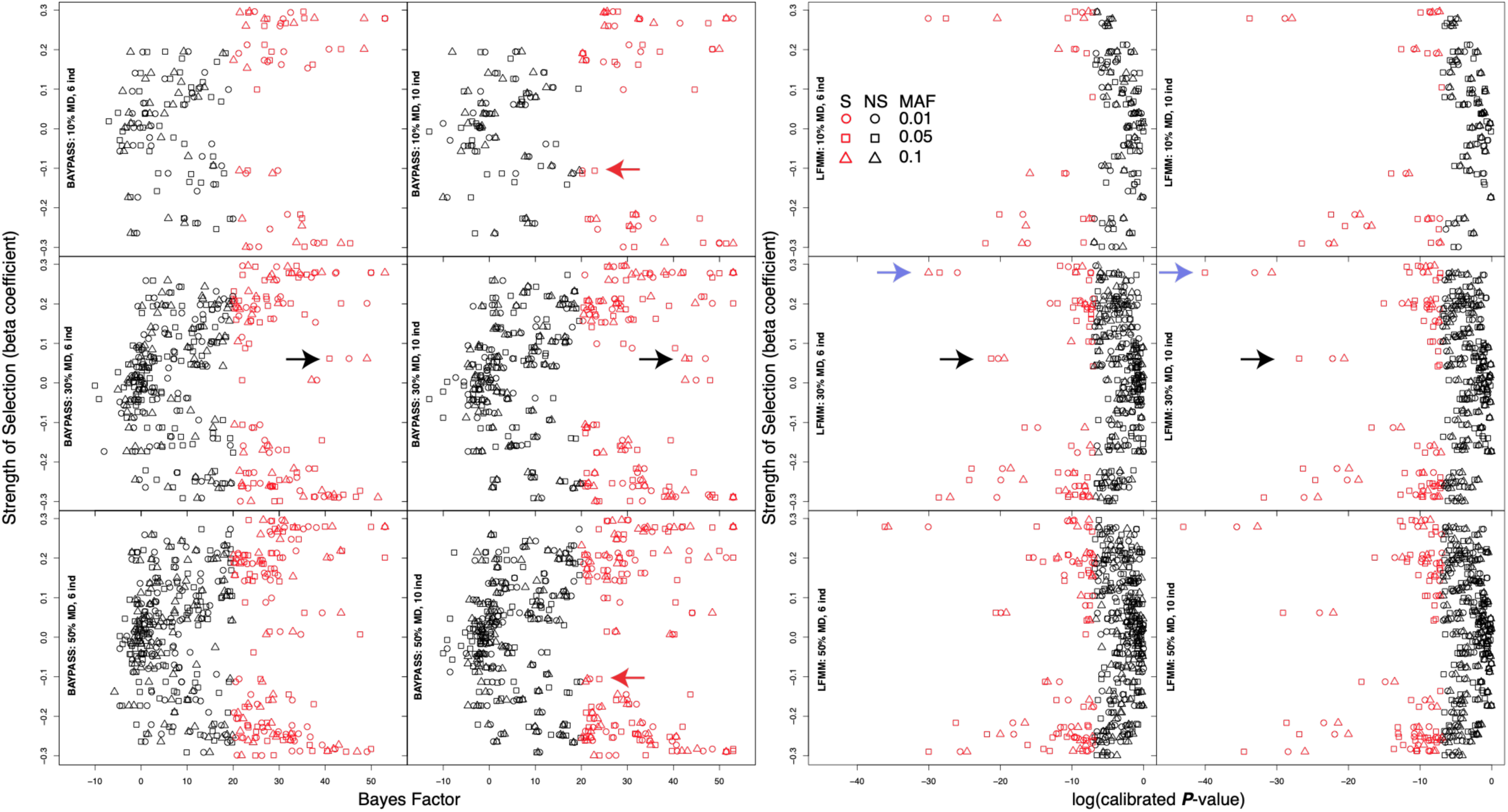
The strength of selection for each SNP and the resulting power of association for BayPass (Bayes Factor) and LFMM2 (calibrated *P*-value) for *Sim calophylla*. S = significant (red); NS = not significant (black). See text for explanations of red, blue and black arrows.

Increasing the strength of selection increased the rate of TP detection. However, false negatives (SNPs under selection not detected as significant) occurred across all selection strengths (Figure 4). The proportion of missing data appeared to have more of an effect on identifying TPs than the number of samples, but this is likely due to differences in the total number of SNPs in the dataset and not due to missing data *per se* (Figure S2). There was little change in the number of TPs identified whether 6 or 10 individuals per population were sampled. Furthermore, as more data were retained through less stringent filtering of missing data, we could identify TPs under weaker selection for both BayPass and LFMM2. However, even with adjustments to sample size and the amount of missing data (number of SNPs retained), a large proportion of TPs were not identified irrespective of filtering parameters. For example, in *Sim calophylla* only 20% (± 3% SD) of the simulated adaptive SNPs were identified by LFMM2, 30% (± 3% SD) by BayPass, and 0% (± 0% SD) by RDA. In analyses of *Sim microcarpa* a higher proportion of the adaptive variants was identified, with 75% (± 5% SD) of the SNPs under selection being identified by LFMM2, 62% (± 3% SD) by BayPass, and 38% (± 2% SD) by RDA.

Minor allele frequency, in combination with biological sample size, impacted the significance of individual SNPs. There were multiple examples where a SNP was considered significant for one MAF but not another (Figure 4 highlights three SNPs indicted by red, black, and blue arrows). One SNP (red arrow, Figure 4) was identified as significant when MAF = 0.01 and 0.05 but not when MAF = 0.1, while holding the number of individuals to 10 and MD at 10%. However, these three SNPs were significant at all MAFs when allowing 50% missing data. Furthermore, the significance of the same SNP with different MAFs can change depending on the method or sample size. For example, one SNP (black arrows, Figure 4) in the Baypass analysis using six individuals per population (162 total), was most significant when MAF = 0.1. When there were 10 individuals per population (270 total) the significance of this SNP was greatest when MAF = 0.01, and lowest when MAF = 0.1. We investigated whether these differences might be due to variation in the covariance (Ω) matrices but found that the covariation among covariance matrices were highly correlated (correlation coefficients ranged between 0.87 and 0.93; all *p*-values < 0.001) and had little effect on the observed differences. One SNP detected in the LFMM2 analyses (blue arrows, Figure 4) showed a significance pattern with MAF 0.1 > 0.05 > 0.01 when there were six individuals per population, but the significance rank changed to MAF 0.05 > 0.01 > 0.1 when there were 10 individuals per population.

While significance levels were significantly (*p* < 0.001) consistent across datasets, LFMM2 had higher consistency with all values >0.98 correlation values while BayPass were between 0.8 and 0.87 for both species (Table S2), slight changes of filtering thresholds did affect outcomes in some circumstances. The influence of MAF on individual SNP significance was observed when comparing significance levels of individual SNPs identified for *Sim calophylla* (Table S3). For BayPass, MAF had a greater effect on the significance level of individual SNPs when using smaller sample sizes (162 vs 270 individuals); more SNPs became non-significant when the biological sample size was smaller. Although the difference in significance level varied with biological sample size, the variation (SD) was similar (Table S3). The opposite was observed with LFMM2 where MAF had less impact (i.e. smaller differences and less variation) on the significance levels of individual SNPs in analyses that used smaller biological sample sizes (Table S3), yet, compared to BayPass, more SNPs became non-significant when changing from MAF = 0.01 to MAF = 0.05.

### Impacts of filtering on extrapolation and interpretation

The impact of filtering the SNP datasets on downstream extrapolation of ‘adaptive’ genomic variation across geographic space varied depending on the thresholds applied and the GEA method. The greatest difference in the climate selection surface (i.e. adaptive predictions through geographic space) between the two approaches and two species was observed for the empirical dataset for *E. microcarpa* using LFMM2 (Figure 5). Applying the conservative thresholds for MAF and MD (while keeping sample size constant) resulted in a significantly different pattern of adaptive genomic variation across the landscape (climate selection surfaces) with different geographic areas predicted to be locally adapted (e.g. red surfaces in Figure 5). This is evident in the comparison between the surfaces produced using the conservative and liberal thresholds using LFMM2 on *E. microcarpa*, where more liberal SNP filtering tended to have a north-south pattern compared to an east-west pattern for the conservative filtering. These contrasting spatial patterns resulted in large differences between predictions (an adaptive index change of > 4). The liberally filtered dataset for LFMM2 was more consistent with both of the predictions for BayPass. Conversely, the effect of filtering was not as apparent when using BayPass on *E. microcarpa*, nor on any of the *C. calophylla* predictions, where, though significant, only subtle differences between filtering thresholds and GEA methods were observed (Figure 5). Nevertheless, the incongruences for *E. microcarpa* occurred along the margins of the species distribution with the conservative filtering method slightly underpredicting putative adaption compared to the liberally filtered dataset. Likewise, incongruences for *C. calophylla* adaptive predictions showed statistically significant differences along the species margins but also within the interior region for both GEA methods, although LFMM2 had slightly larger incongruences compared to BayPass. Consistent with the empirical datasets, the general spatial patterns of adaptive variation predicted using the simulated datasets remained qualitatively the same despite filtering for MAF, MD, and sample size, indicating that the signal to noise remained quite similar despite the higher number of FPs in the more liberal datasets (Figure S4). However, we detected regions where there were statistically significant differences among datasets, particularly along the margins of the species’ ranges. Those differences were driven by different filtering parameters and GEA methods. For instance, the biggest changes for *Sim microcarpa* in BayPass are driven by MAF, but missing data and number of samples had the biggest impact in LFMM2 (Figure S4). The increase in the number of individuals from 260 total individuals to 650 individuals had very little impact on landscape-wide patterns of genomic variation (i.e. adaptive index).

**Figure 5.**
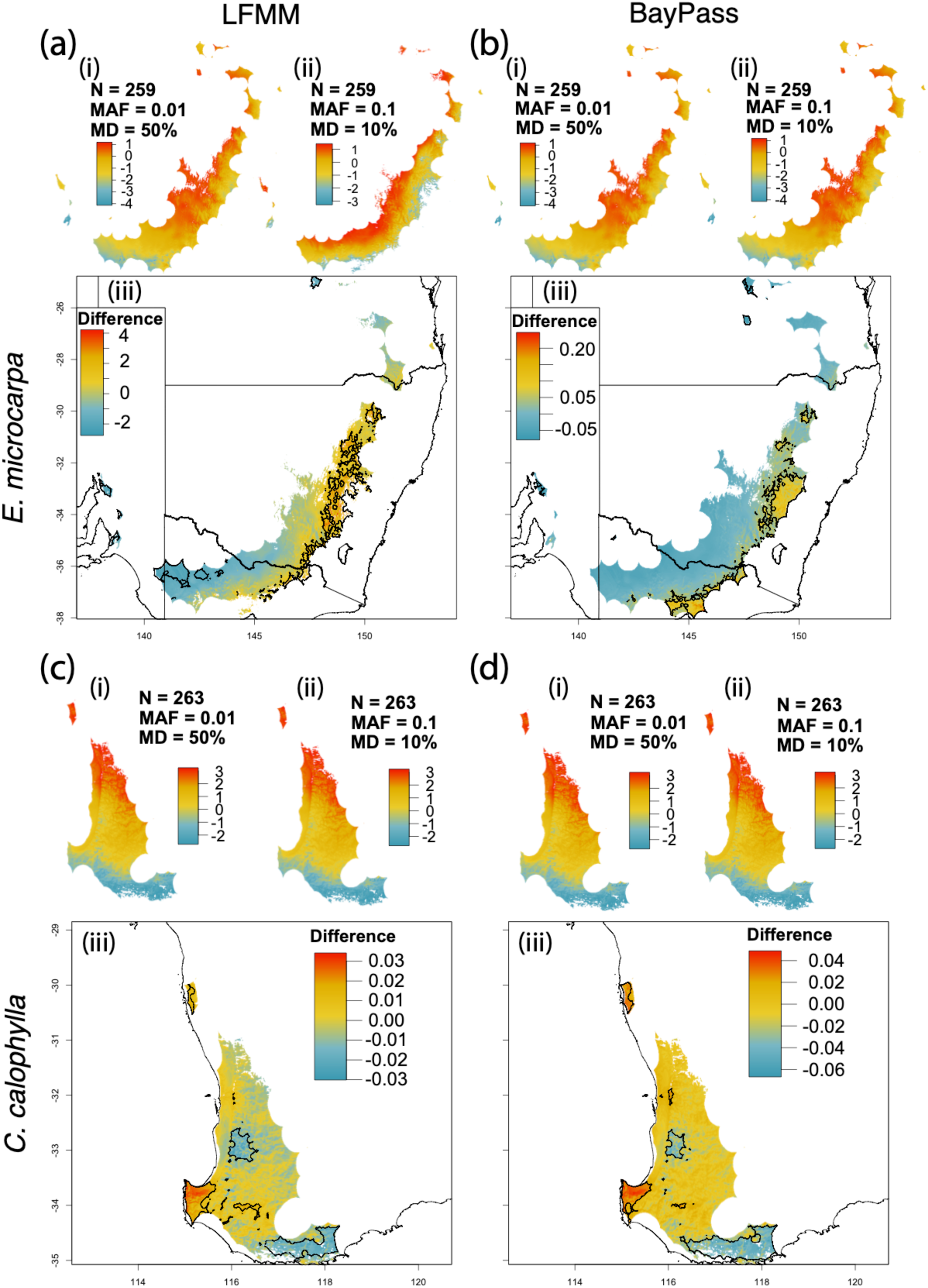
Analysis of the effect of filtering on spatial extrapolation of adaptive variation within the empirical datasets. The maps within boxes (iii) show the differences between the ‘liberal’ (i) and ‘conservative’ (ii) maps (smaller maps directly above). Combinations of species are *E. microcarpa* and LFMM2 (a), *E. microcarpa* and Baypass (b), *C. calophylla* and LFMM2 (c), and *C. calophylla* and BayPass (d). Red surface colours in the smaller maps represent regions of each species gene pool putatively adapted to hotter and drier climates while the blue surface represents the regions putatively adapted to increasingly cooler and wetter climates. The red surface in the main differential maps represent regions where the liberal dataset predicted stronger adaptation, whereas the blue surface corresponds to regions where the conservative dataset predicted stronger adaptation. Note: the differential scales are different across comparisons, this was done to highlight the differences within each comparison. Areas of significant differences in predicted magnitude of adaptation are outlined with a black polygon. Liberal dataset = missing data (MD) = 50%; minor allele frequency (MAF) = 0.01. Conservative dataset = MD = 10%; MAF = 0.1; MAF = minor allele frequency; MD = missing data; N = sample size.

## Discussion

Most studies filter data prior to GEA analysis with the aim of improving the quality of the input data to obtain better inferences of environmental adaptation. While several studies have explored the influence of demographic history, population structure, sampling strategy, landscape configuration, and strength of selection on the capacity of various approaches to detect loci under selection (Forester et al., 2018; Lotterhos & Whitlock, 2014, 2015; Luu, Bazin, & Blum, 2016; de Mita et al., 2013; Rellstab, Gugerli, Eckert, Hancock, & Holderegger, 2015; Schlamp et al., 2015; de Villemereuil et al., 2014), the impact of filtering thresholds on GEA outputs has not been thoroughly evaluated previously. Given the wide range of filtering choices used and the lack of broad scale patterns of adaptation (Ahrens et al., 2018), we explored the impact of filtering on the capacity of various approaches to identify putatively adaptive SNPs, and demonstrated that filtering thresholds do impact the outcomes of GEA analyses. We reveal that filtering for minor allele frequency and missing data affects GEA outputs in various ways depending upon species, sample size, and GEA analytical method. To summarise how our study challenges the four common assumptions addressed in the introduction:

1. More stringent filtering reduces the identification of FPs but the rate of identifying FPs remains constant across most filtering thresholds.
2. Loci with strong selection strengths are more likely to be identified as TPs but a strong selection strength does not guarantee a significant identification.
3. Combining GEA analyses slightly reduces FPs but at the expense of TPs.
4. Predictions across the landscape, for the most part, were biologically robust but statistically different across all filtering thresholds, although some circumstances led to biologically and statistically different adaptive patterns.

Ultimately, we found that filtering choices can have multiplicative effects for downstream interpretation, meaning that small filtering changes could change estimates of genomic predicted adaptation to the environment. While we focused on widespread tree species, the concepts drawn from our results are applicable across other organisms and we suggest that some common practices employed in GEA studies should be reconsidered.

### Effects of filtering on GEA outputs

Missing data are usually minimised in order to improve the reliability of the dataset. However, our results suggest that filtering data with strict missing data thresholds does not necessarily improve GEA outcomes. In fact, filtering missing data seemed to have little effect on the ratio of TP to AA. This is in line with other population genetic studies that found that missing data (within reason) do not affect calculations of *F*_ST_ or H_e_ (Binks, Gibson, Ottewell, Macdonald, & Byrne, 2019; Díaz-Arce & Rodríguez-Ezpeleta, 2019; Shafer et al., 2017). Indeed, we found that BayPass, LFMM2, and RDA (specific to *Sim microcarpa*) were robust to missing data with respect to TP:AA but the actual number of TPs and FPs identified varied. For BayPass and RDA, this could be partially due to the use of population-level allele counts or allele frequencies as the input data, a strategy that effectively ignores missing data. Because LFMM2 uses individual genotypes, and we naively imputed the gaps using loci means (default parameter), we expected that missing data would result in more FPs and thereby provide a possible source of differentiation among methods (de Villemereuil et al., 2014). While this was apparent for *Sim calophylla*, LFMM2 performed well for *Sim microcarpa*. The ‘missingness’ was similar among species, meaning that the different responses between species suggest that the relatively high FP call rate for *Sim calophylla* is likely due to a combination of missingness and other underlying differences between the species.

The lack of improvement to GEA outputs with decreasing proportions of missing data suggests that the number of SNPs in a dataset is more important than dataset completeness, within reason, bearing in mind that we only tested up to 50% missing data. More SNPs allow sufficiently large numbers to statistically define ‘neutral demographic structure’, an important aspect to all GEA analyses, and thus increase the number of putatively adaptive SNPs identified (see further discussion below). The relative importance placed on filtering missing data should depend on the downstream application of putatively adaptive loci. This is borne out by the maps in Figure S4 (particularly between the missing data thresholds), where the presence of more FPs do not affect the adaptive signal to non-adaptive noise, at least when the signal from TPs is sufficiently large. However, this interpretation must be qualified, because the discovery of TPs in empirical datasets is unknown, and it is the strength and number of TPs that will override a contrasting FP signal.

Minor allele frequency is an important threshold, because nonsynonymous SNPs are likely to have a MAF less than 0.05 (Cargill et al., 1999) and, in human studies, inclusion of SNPs with low MAF increases the rate of identification of causal variants (Gorlov, Gorlova, Sunyaev, Spitz, & Amos, 2008). Our data suggest that a low minor allele frequency has a type I error (FP) rate close to nominal levels (i.e. FP rate is similar among datasets), which has been found in other studies (Moskvina, Craddock, Holmans, Owen, & O’Donovan, 2006; Tabangin, Woo, & Martin, 2009). These findings suggest that low MAF should not be excluded from GEA datasets if sampling design is sufficiently large. However, in our study, MAF influenced FP call rates with varying impacts between programs and species. It is important to note that MAF filtering is also a function of sample size and missing data. The larger the sample size, the smaller the MAF threshold can be. This is most apparent when considering MAF as minor allele counts (MAC; see O’Leary et al., 2018 for discussion), where a low MAF could still result in a high MAC for larger sample sizes, allowing for sequencing error issues to be resolved by maintaining SNPs that are called confidently (higher MAC). Ultimately, like missing data, MAF affects the total number of SNPs in the dataset, but it can also influence a SNP’s significance.

While more stringent filtering may, theoretically, improve the quality of the dataset, the reduction in the overall size of the data set and the potential loss of informative loci may influence the null models underlying GEA analyses and thus the identification of SNP-environment associations. This is evident in the impact of both missing data and MAF on the detection of adaptive SNPs under different strengths of selection. Both Baypass and LFMM2 missed TPs at all selection strengths, even for SNPs under strong selection pressure. However, datasets that included more missing data yielded TPs that were under weak selection (∼0.05). This is likely because less stringent filtering of missing data results in larger datasets, thereby increasing the overall number of TPs. In addition, despite the missing data, larger datasets (relative to reduced representation datasets with 2-20k SNPs) may enable a more statistically significant ‘null model’ for the GEA and therefore greater power to detect loci under selection (Morin, Martien, & Taylor, 2009); and the power of the number of SNPs in genome-wide association studies has been discussed previously (Hong & Park, 2012; Klein, 2007; Spencer, Su, Donnelly, & Marchini, 2009), and the same logic applies for GEAs. Filtering of MAF may also influence the null model, changing the significance of TPs and, thus, their potential to be identified as TPs. While stringently filtering genomic data may create a more reliable dataset in theory, having fewer data points appears to reduce the overall power and effectiveness of GEAs.

### Combining results

Using loci identified across multiple analyses reduced both the number of FPs and TPs. This is common practice and in one sense, our results support this commonly-used approach (Forester et al., 2018; Lotterhos & Whitlock, 2015) in that we observed a slight increase in TP:AA. However, the TPs retained reflected the more conservative analysis, and most of the TPs identified by the other methods were lost. Each method uses unique approaches to identify SNPs (e.g. controlling for population structure and statistical model) and different methods are likely to identify different suites of putatively adaptive SNPs. This output agrees with findings from Forester et al., (2018); that combining results will bias the results to strong selective sweeps and limit findings to the least powerful (or most conservative) method. The trade-off between reduced FPs and the loss of informative TPs therefore needs careful consideration, particularly given that downstream extrapolation of results tends to be largely unaffected by the presence of FPs. If one uses an overlapping approach we suggest using the Lotterhos et al. (2017) composite measure to improve the identification of adaptive signals by using the outputs across many GEA methods.

### Influence on downstream applications

For the most part, the patterns of geospatial predictions were biologically similar but statistically different within species and methods, but across filtering thresholds. However, this was not the case for *E. microcarpa* and LFMM2. The difference between the liberal and conservative datasets revealed different biological and statistical geospatial adaptive patterns. The more liberal dataset was more similar to both BayPass outputs, suggesting that the LFMM2 conservative prediction was spurious. While it is possible that both patterns are correct due to hierarchically complex relationships between adaptation and climate, this pattern is likely due to the fact that the FPs had a larger impact on the adaptive signal because there were fewer TPs overall (i.e. the noise was greater than the signal), as only five putatively adaptive SNPs associated with BIO5 (eight for BIO14) were identified in the conservative dataset compared to 101 putatively adaptive SNPs associated with BIO5 (36 for BIO14) in the liberal dataset. This outcome suggests that FPs can affect predictions when fewer TPs are found for LFMM2, but this effect was lost when more TPs are kept through larger datasets and liberal filters.

Pseudo positives (PP) were found to be a confounding factor, particularly for the *Sim calophylla* dataset. Indeed, the correlation coefficients of the two environmental variables suggested that PPs would have a much greater impact on the *Sim calophylla* dataset than on the *Sim microcarpa* dataset. While we did find PPs in *Sim calophylla*, they numbered only about 20% of the number of TPs; this was less than expected considering the strong correlation of the two climatic variables across the distribution of *C. calophylla*. This suggests that it is preferable to include environmental variables that are not correlated in GEA analyses (see Hoban et al., 2016); however, the inclusion of variables with correlation coefficients around 0.7 seems to be adequate (agreeing with the findings in Dormann et al., (2013)), particularly if they were chosen a priori with hypothesis-driven questions.

In our simulations, we chose higher than expected strength-of-selection coefficients to try to identify selection coefficients that would enable identification of adaptive SNPs above a given threshold. We were unable to identify a consistent threshold and therefore conclude that strong selection pressure is not sufficient to identify adaptive SNPs, and that the SNPs must be distributed throughout the populations in specific ways. However, we did find a strong relationship between the significance of SNPs and strength-of-selection, indicating that, not surprisingly, there is a much higher probability of identifying SNPs of large effect using either of the univariate methods than with RDA.

### Differences among species

The datasets that we examined showed different responses to the effects of filtering despite being (i) derived from related species that span similar climate gradients, and (ii) produced using the same reduced representation approach. One reason for these differences could be the different genome sizes of these species. The genome of *C. calophylla* is estimated to be 400 Mb while that of *E. microcarpa* is around 700 Mb. Although genome size is likely not evolutionarily significant (Vu et al., 2015), it could influence the search for adaptive SNPs, as a smaller genome size would provide better representation of coding regions. A second reason for the differences between datasets could be that, even though the two species inhabit similar temperature gradients, the broader climate of each species is fundamentally different: *C. calophylla* occurs in a Mediterranean-type climate and *E. microcarpa* occurs in a temperate climate. A third reason for the differences between species could be that similar levels of global population structure does not dictate how genetic variance is distributed within species. For instance, it is possible that when more SNPs are kept due to filtering thresholds, the estimated population structure may change in different ways for each species. Finally, species’ geographic range size, as well as demographic and evolutionary history, may explain differences in results. *Eucalyptus microcarpa* has a larger geographic range than *C. calophylla*, indicating that underlying demographic history could be fundamentally different (e.g. expansion/contraction).

### Conclusions

While we provide a filtering roadmap that enables users to understand how filtering might affect GEA outputs, all organisms and datasets we study are unique, and the questions developed for each will be different. Therefore, there is no universally ‘best’ way to perform filtering for GEA analyses. Datasets should be developed in ways that best fit the objectives of the study (some possible examples and recommendations are given in Table 4). Another important component that we have not addressed, and is outside the scope of this study, is the use of genomic resources for the betterment of GEA outputs. Additional genomic resources, such as an annotated reference genome, provide further chances to refine the SNP sets used for downstream analyses or applications. For example, it might be useful to examine whether SNPs that putatively mediate local adaptation are located near genes whose function is relevant to the environmental variable (Manel et al., 2016), or whose expression is induced by relevant environmental challenges. Collectively, if a large proportion of putatively adaptive SNPs are located near genes with relevant functions, it might promote confidence in the associations, and their application to management actions.

**Table 4.** Outcomes and suggestions of different filtering approaches for different project aims employing GEA analyses. TP = True Positive; FP = False Positive; MD = missing data; MAF = minor allele frequency; MAC = minor allele count.

| Aim / Concern | Example research question or application | Filtering approach | Analysis outcome |
| --- | --- | --- | --- |
| Conservation | Understanding general landscape patterns of genomic diversity for conservation or management. Reference genome may not be available. | More relaxed filtering to create larger overall SNP dataset. Smaller permissible MAF (given sample size and thus MAC). Larger amount of MD. Pool unique candidate SNPs across multiple methods. | Large overall pool of 'adaptive' SNPs, including mix of TPs and FPs; providing overview of the adaptive landscape. |
| Maximise TP | Patterns of genomic adaptation across major environmental gradients. Reference genome available. | More relaxed filtering. Larger amount of MD. Smaller permissible MAF. Pool candidate SNPs from multiple methods (don't just select overlapping results). Refine SNP sets with location and/or functional annotation. | Larger overall dataset of 'adaptive' SNPs; maximising number of TPs and improving dataset for downstream applications. |
| Minimise FP | Looking for candidate large-effect loci under selection for further investigation, especially where no genome is available. | More stringent filtering. Fewer MD. Consider MAF as a function of sample size and missing data (MAC) as well as impacts on significance. Focus on SNPs occurring in multiple programs using the Lotterhos et al., (2017) composite method. | Decreased absolute number of FPs at the expense of the number of TPs. Reduced identification of loci under weaker selection. |
| Identify loci under weak selection | Quantification of genome-wide levels of adaptation driven by environmental selection. Reference genome may or may not be available. | More relaxed filtering. Larger amount of MD. Lower permissible MAF with larger biological sample sizes. Refine SNP sets with location and/or functional annotation. | Increased power of GEA analyses. Greater number of loci providing more informative null models for GEA analyses. Improved ability to detect loci under weaker selection. |

Identifying true adaptive variants is difficult, particularly for non-model organisms, and this is true even when strengths-of-selection are large. When we try to create and use the most complete datasets through stringent filtering, we filter out many of those strongly adaptive SNPs that are likely to be identified as TPs. When we have fewer putatively adaptive SNPs, then the noise of FPs might lead to spurious adaptive signals through predictions, as we show. On the other hand, if we filter our datasets more liberally, the adaptive signal seems to overpower spurious signals. Together, as we identify clearer signals of adaptation, we are likely to better understand how non-model species have adapted to the environment, moving the field of landscape genomics toward a more complete understanding of our natural systems.

## Data accessibility

All data will be uploaded to dryad upon acceptance and R code will be made available through github or dryad.

## Author contributions

CA developed the original idea. All authors contributed to further development of the idea at a workshop hosted at Western Sydney University. CA, TH, KM, PH, RA, and JB developed the code and analytics pipeline. CA and RJ wrote the first draft. All authors edited various versions of the manuscript.

## Acknowledgements

We acknowledge Prof James Seeb’s use of “F-word” at the CONGEN 2013 meeting which is discussed in Andrews & Luikart (2014) manuscript. This paper gained traction at the Eucalyptus Australia meeting in 2019 in Hobart, where there was interest to compare adaptive patterns across eucalypt species, and therefore thank the organisers and presenters at the meeting for the opportunity to meet and discuss the genomics of eucalypts.

## References

Ahrens, C. W., Byrne, M., & Rymer, P. D. (2019). Standing genomic variation within coding and regulatory regions contributes to the adaptive capacity to climate in a foundation tree species. Molecular Ecology, 28(10), 2502–2516.

Ahrens, C. W., Rymer, P. D., Stow, A., Bragg, J., Dillon, S., Umbers, K. D. L., & Dudaniec, R. Y. (2018). The search for loci under selection: trends, biases and progress. Molecular Ecology, 27(6), 1342–1356.

Ahrens, C.W., James, E.A., Miller, A.D., Ferguson, S., Aitken, N.C., Jones, A.W., Lu-Irving, P., Borevitz, J.O., Cantrill, D.J. and Rymer, P.D. (2020). Spatial, climate, and ploidy factors drive genomic diversity and resilience in the widespread grass *Themeda triandra*. Molecular Ecology, doi: 10.1111/mec.15614

[dataset] Ahrens, C.W., Jordan, R., Bragg, J., Harrison, P.A., Hopley, T., Bothwell, H.,… (2020). Regarding the F-word: the effects of data Filtering on inferred genotype-environment associations. DOI: (to be provided upon acceptance via dryad – data and R code)

Andrews, K. R., & Luikart, G. (2014). Recent novel approaches for population genomics data analysis. Molecular Ecology, 23(7), 1661–1667.

Bay, R. A., Harrigan, R. J., Le Underwood, V., Gibbs, H. L., Smith, T. B., & Ruegg, K. (2018). Genomic signals of selection predict climate-driven population declines in a migratory bird. Science, 359(6371), 83–86.

Binks, R. M., Gibson, N., Ottewell, K. M., Macdonald, B., & Byrne, M. (2019). Predicting contemporary range-wide genomic variation using climatic, phylogeographic and morphological knowledge in an ancient, unglaciated landscape. Journal of Biogeography, 46(3), 503–514.

Browne, L., Wright, J. W., Fitz-Gibbon, S., Gugger, P. F., & Sork, V. L. (2019). Adaptational lag to temperature in valley oak (*Quercus lobata*) can be mitigated by genome-informed assisted gene flow. Proceedings of the National Academy of Sciences, 116(50), 25179–25185.

Cargill, M., Altshuler, D., Ireland, J., Sklar, P., Ardlie, K., Patil, N., … Lander, E. S. (1999). Characterization of single-nucleotide polymorphisms in coding regions of human genes. Nature Genetics, 22(3), 231–238.

Caye, K., Jumentier, B., Lepeule, J., & François, O. (2019). LFMM 2: Fast and Accurate Inference of Gene-Environment Associations in Genome-Wide Studies. Molecular Biology and Evolution, 36(4), 852–860.

Coop, G., Witonsky, D., Rienzo, A. D., & Pritchard, J. K. (2010). Using environmental correlations to identify loci underlying local adaptation. Genetics, 185(4), 1411–1423.

Costa e Silva, J., Potts, B., Harrison, P. A., & Bailey, T. (2019). Temperature and rainfall are separate agents of selection shaping population differentiation in a forest tree. Forests, 10(12), 1145.

Díaz-Arce, N., & Rodríguez-Ezpeleta, N. (2019). Selecting RAD-Seq data analysis parameters for population genetics: the more the better? Frontiers in Genetics, 10, 533.

Dormann CF, Elith J, Bacher S, Buchmann C, Carl G, Carré G, Marquéz JRG, Gruber B, Lafourcade B, Leitão PJ, Münkemüller T, McClean C, Osborne PE, Reineking B, Schröder B, Skidmore AK, Zurell D, Lautenbach S (2013) Collinearity: a review of methods to deal with it and a simulation study evaluating their performance. Ecography, 36, 27–46.

Fick, S. E., & Hijmans, R. J. (2017). WorldClim 2: new 1-km spatial resolution climate surfaces for global land areas. International Journal of Climatology, 37(12), 4302–4315.

Forester, B. R., Lasky, J. R., Wagner, H. H., & Urban, D. L. (2018). Comparing methods for detecting multilocus adaptation with multivariate genotype–environment associations. Molecular Ecology, 27(9), 2215–2233.

François, O., Martins, H., Caye, K., & Schoville, S. D. (2016). Controlling false discoveries in genome scans for selection. Molecular Ecology, 25(2), 454–469.

Fu, Y.-B. (2014). Genetic diversity analysis of highly incomplete SNP genotype data with imputations: an empirical assessment. G3: Genes|Genomes|Genetics, 4(5), 891–900.

Garner, B. A., Hand, B. K., Amish, S. J., Bernatchez, L., Foster, J. T., Miller, K. M., … Luikart, G. (2016). Genomics in conservation: case studies and bridging the gap between data and application. Trends in Ecology & Evolution, 31(2), 81–83.

Gautier, M. (2015). Genome-wide scan for adaptive divergence and association with population-specific covariates. Genetics, 201(4), 1555–1579.

Gautier, M., Gharbi, K., Cezard, T., Foucaud, J., Kerdelhué, C., Pudlo, P., … Estoup, A. (2012). The effect of RAD allele dropout on the estimation of genetic variation within and between populations. Molecular Ecology, 22(11), 3165–3178.

Gorlov, I. P., Gorlova, O. Y., Sunyaev, S. R., Spitz, M. R., & Amos, C. I. (2008). Shifting paradigm of association studies: value of rare single-nucleotide polymorphisms. The American Journal of Human Genetics, 82(1), 100–112.

Günther, T., & Coop, G. (2013). Robust identification of local adaptation from allele frequencies. Genetics, 195(1), 205–220.

Hendricks, S., Anderson, E. C., Antao, T., Bernatchez, L., Forester, B. R., Garner, B., … Luikart, G. (2018). Recent advances in conservation and population genomics data analysis. Evolutionary Applications, 11(8), 1197–1211.

Hoban, S., Kelley, J. L., Lotterhos, K. E., Antolin, M. F., Bradburd, G., Lowry, D. B., … Whitlock, M. C. (2016). Finding the genomic basis of local adaptation: pitfalls, practical solutions, and future directions. The American Naturalist, 188(4), 379–397.

Hong, E. P., & Park, J. W. (2012). Sample size and statistical power calculation in genetic association studies. Genomics & Informatics, 10(2), 117–122.

Jeffreys, H. (1961). Theory of probability, 3rd Edn Oxford: Oxford University Press. Oxford, UK.

Jordan, R., Hoffmann, A. A., Dillon, S. K., & Prober, S. M. (2017). Evidence of genomic adaptation to climate in *Eucalyptus microcarpa*: implications for adaptive potential to projected climate change. Molecular Ecology, 26(21), 6002–6020.

Klein, R. J. (2007). Power analysis for genome-wide association studies. BMC Genetics, 8(1), 58.

Linck, E., & Battey, C. J. (2019). Minor allele frequency thresholds strongly affect population structure inference with genomic data sets. Molecular Ecology Resources, 19(3), 639–647.

Lotterhos, K. E., Card, D. C., Schaal, S. M., Wang, L., Collins, C., & Verity, B. (2017). Composite measures of selection can improve the signal-to-noise ratio in genome scans. Methods in Ecology and Evolution, 8(6), 717–727.

Lotterhos, K. E., & Whitlock, M. C. (2014). Evaluation of demographic history and neutral parameterization on the performance of FST outlier tests. Molecular Ecology, 23(9), 2178–2192. doi: 10.1111/mec.12725

Lotterhos, K. E., & Whitlock, M. C. (2015). The relative power of genome scans to detect local adaptation depends on sampling design and statistical method. Molecular Ecology, 24(5), 1031–1046.

Lowry, D. B., Hoban, S., Kelley, J. L., Lotterhos, K. E., Reed, L. K., Antolin, M. F., & Storfer, A. (2017). Breaking RAD: an evaluation of the utility of restriction site-associated DNA sequencing for genome scans of adaptation. Molecular Ecology Resources, 17(2), 142–152.

Luu, K., Bazin, E., & Blum, M. G. B. (2016). pcadapt?: an R package to perform genome scans for selection based on principal component analysis. Molecular Ecology Resources, 17(1), 67–77.

Manel, S., Perrier, C., Pratlong, M., Abi-Rached, L., Paganini, J., Pontarotti, P., & Aurelle, D. (2016). Genomic resources and their influence on the detection of the signal of positive selection in genome scans. Molecular Ecology, 25(1), 170–184.

Mastretta-Yanes, A., Arrigo, N., Alvarez, N., Jorgensen, T. H., Piñero, D., & Emerson, B. C. (2015). Restriction site-associated DNA sequencing, genotyping error estimation and de novo assembly optimization for population genetic inference. Molecular Ecology Resources, 15(1), 28–41.

Meirmans, P. G. (2015). Seven common mistakes in population genetics and how to avoid them. Molecular Ecology, 24(13), 3223–3231.

de Mita, S., Thuillet, A., Gay, L., Ahmadi, N., Manel, S., Ronfort, J., & Vigouroux, Y. (2013). Detecting selection along environmental gradients: analysis of eight methods and their effectiveness for outbreeding and selfing populations. Molecular Ecology, 22(5), 1383–1399.

Morin, P. A., Martien, K. K., & Taylor, B. L. (2009). Assessing statistical power of SNPs for population structure and conservation studies. Molecular Ecology Resources, 9(1), 66–73.

Moskvina, V., Craddock, N., Holmans, P., Owen, M. J., & O’Donovan, M. C. (2006). Effects of differential genotyping error rate on the type I error probability of case-control studies. Human Heredity, 61(1), 55–64.

Nadeau, S., Meirmans, P. G., Aitken, S. N., Ritland, K., & Isabel, N. (2016). The challenge of separating signatures of local adaptation from those of isolation by distance and colonization history: the case of two white pines. Ecology and Evolution, 6(24), 8649–8664.

Narum, S. R., Buerkle, C. A., Davey, J. W., Miller, M. R., & Hohenlohe, P. A. (2013). Genotyping-by-sequencing in ecological and conservation genomics. Molecular Ecology, 22(11), 2841–2847.

Nazareno, A. G., Bemmels, J. B., Dick, C. W., & Lohmann, L. G. (2017). Minimum sample sizes for population genomics: an empirical study from an Amazonian plant species. Molecular Ecology Resources, 17(6), 1136–1147.

O’Leary, S. J., Puritz, J. B., Willis, S. C., Hollenbeck, C. M., & Portnoy, D. S. (2018). These aren’t the loci you’e looking for: Principles of effective SNP filtering for molecular ecologists. Molecular Ecology, 27(16), 3193–3206.

Orsini, L., Mergeay, J., Vanoverbeke, J., & Meester, L. (2013). The role of selection in driving landscape genomic structure of the waterflea *Daphnia magna*. Molecular Ecology, 22(3), 583–601.

Pool, J. E., Hellmann, I., Jensen, J. D., & Nielsen, R. (2010). Population genetic inference from genomic sequence variation. Genome Research, 20(3), 291–300.

Prober, S. M., Potts, B. M., Bailey, T., Byrne, M., Dillon, S., Harrison, P. A., … Vaillancourt, R. E. (2016). Climate adaptation and ecological restoration in eucalypts. Proceedings of the Royal Society of Victoria, 128(1), 40.

Razgour, O., Forester, B., Taggart, J. B., Bekaert, M., Juste, J., Ibáñez, C., … Manel, S. (2019). Considering adaptive genetic variation in climate change vulnerability assessment reduces species range loss projections. Proceedings of the National Academy of Sciences, 116(21), 201820663.

Rellstab, C., Gugerli, F., Eckert, A. J., Hancock, A. M., & Holderegger, R. (2015). A practical guide to environmental association analysis in landscape genomics. Molecular Ecology, 24(17), 4348–4370.

Schlamp, F., Made, J. van der, Stambler, R., Chesebrough, L., Boyko, A. R., & Messer, P. W. (2015). Evaluating the performance of selection scans to detect selective sweeps in domestic dogs. Molecular Ecology, 25(1), 342–356.

Sgrò, C. M., Lowe, A. J., & Hoffmann, A. A. (2011). Building evolutionary resilience for conserving biodiversity under climate change. Evolutionary Applications, 4(2), 326–337.

Shafer, A. B. A., Peart, C. R., Tusso, S., Maayan, I., Brelsford, A., Wheat, C. W., & Wolf, J. B. W. (2017). Bioinformatic processing of RAD-seq data dramatically impacts downstream population genetic inference. Methods in Ecology and Evolution, 8(8), 907–917.

Sork, V. L., Aitken, S. N., Dyer, R. J., Eckert, A. J., Legendre, P., & Neale, D. B. (2013). Putting the landscape into the genomics of trees: approaches for understanding local adaptation and population responses to changing climate. Tree Genetics & Genomes, 9(4), 901–911.

Sork, Victoria L. (2017). Genomic studies of local adaptation in natural plant populations. Journal of Heredity, 109(1), 3–15.

Spencer, C. C. A., Su, Z., Donnelly, P., & Marchini, J. (2009). Designing genome-wide association studies: sample size, power, imputation, and the choice of genotyping chip. PLoS Genetics, 5(5), e1000477.

Storz, J. F. (2005). Using genome scans of DNA polymorphism to infer adaptive population divergence. Molecular Ecology, 14(3), 671–688.

Tabangin, M. E., Woo, J. G., & Martin, L. J. (2009). The effect of minor allele frequency on the likelihood of obtaining false positives. BMC Proceedings, 3(Suppl 7), S41.

Thornhill, A. H., Crisp, M. D., Külheim, C., Lam, K. E., Nelson, L. A., Yeates, D. K., & Miller, J. T. (2019). A dated molecular perspective of eucalypt taxonomy, evolution and diversification. Australian Systematic Botany, 32(1), 29–48.

Vu, G. T. H., Schmutzer, T., Bull, F., Cao, H. X., Fuchs, J., Tran, T. D., … Schubert, I. (2015). Comparative genome analysis reveals divergent genome size evolution in a carnivorous plant genus. The Plant Genome, 8(3), 1–14.

Willing, E.-M., Dreyer, C., & Oosterhout, C. van. (2012). Estimates of genetic differentiation measured by FST do not necessarily require large sample sizes when using many SNP markers. PLoS ONE, 7(8), e42649.

Zuur, A. F., Ieno, E. N., & Elphick, C. S. (2010). A protocol for data exploration to avoid common statistical problems. Methods in ecology and evolution, 1(1), 3–14.

