## Supplementary material for "Regarding the *F*-word: the effects of data *Filtering* on inferred genotype-environment associations"

Table of contents


| Table/Figure | Description | Page # |
|---|---|---|
| Methods | | 2 |
| Results | | 3-4 |
| Table S1 | Spatial autocorrelation and effective sample size calculations for all climate variables. | 5 |
| Table S2 | Correlation between significant values. | 6 |
| Table S3 | Changes of significance for SNPs between the different MAFs. | 7 |
| Figure S1 | Distribution of missing data in simulated datasets. | 8 |
| Figure S2 | The relationship between total number of SNPs and the number of associations. | 9 |
| Figure S3 | The relationship between strength of selection and significance. | 10 |
| Figure S4 | Effect of minor allele frequency, missing data, and sample size on adaptive predictions. | 11 |

**Methods**

Assessments of spatial autocorrelation and effective population size

Spatial dependence in the form of spatial autocorrelation is common among environmental variables. Spatial autocorrelation can negatively impact the power of GEA analyses (Manel et al. 2012) by reducing the effective sample size for analysis, while also inflating significance if samples are not independent. Therefore, we explored spatial autocorrelation and $n_{\text{eff-env}}$ of each species for both environmental variables, using the *Moran.I* function in the *spdep* R package (Bivand 2019). We then estimated the $n_{\text{eff-env}}$ following Griffith (2005), taking into consideration the sampling design and spatial autocorrelation of the climate variable. Furthermore, to understand the potential for confounding results we estimated the correlation between the two climate variables (BIO5 & BIO14) using Pearson's correlation coefficient (*r*) among populations within each species, using the *cor* function in R.

Impacts of filtering on extrapolation and interpretation of adaptive variation

This was achieved using the *rda* function of *vegan* in R, which first defines the adaptively-enriched genomic space using the identified climate-associated SNPs and then identifies the linear combination of standardised climate variables (we use the results from both BIO5 and BIO14), which best explains this adaptively-enriched genomics space ('adaptive index' from here on). The canonical coefficients for the first constrained RDA axis were used to map the 'climate selection surface' using the *raster* package of R. This climate selection surface was clipped to a convex hull around each species' native distribution. To determine whether we recover the same adaptive index across the different filtering methods, we performed a Spearmans rank correlation. Spatial differences between the climate selection surface created using each filtered dataset were determined using a pixel pairwise z-score test, executed with the *SigDiff* function of *SDMTools* in R (VanDerWal, Falconi, Januchowski, Shoo, & Storlie, 2019).

**Results**

We used Moran's *I* to provide an indication of the relative power of the sampling design, given the spatial autocorrelation inherent in the selected climate variables. We detected the presence of spatial autocorrelation in both datasets for both climate variables Moran's *I* for BIO5 was 0.33 ($p$ = 1.14e-09) for *C. calophylla* and 0.27 ($p$ = 9.10e-15) for *E. microcarpa* (Table 1). Therefore, not all sampled populations were independent, essentially creating pseudoreplicates within the dataset and decreasing the power with which to test for environmental associations. Similar values of *I* for BIO5 for both species suggests similar effect of spatial autocorrelation on analysis for both species (Table 1), although *I* was slightly greater for *C. calophylla*, leading to low effective sample sizes ($n_{\text{eff-env}}$) for both species: $n_{\text{eff-env}}$ for BIO5 was 15.12 ($p = 1.14e^{-9}$) for *E. microcarpa* and 13.61 ($p = 9.10e^{-15}$) for *C. calophylla*, about half of the sampled number of populations (Table 1). BIO5 and BIO14 were weakly correlated in *E. microcarpa* ($r$ = 0.014; $p$ = 0.95), but negatively correlated in *C. calophylla* ($r$ = -0.75; p < 0.001) (Table 1). For comparisons, we provide the *I* and $n_{\text{eff-env}}$ for all 19 bioclim variables in supplementary information and the results vary broadly with an $n_{\text{eff-env}}$ as low as 11.4 or as high as 23.4 for *E. microcarpa* and 12.5 to 22.4 for *C. calophylla* (Table S1).

Results of sample size and pseudo positives

Larger biological sample sizes consistently identified more TPs for *Sim microcarpa* data than when fewer individuals were included in the dataset (Figure 2f-j). In the *Sim calophylla* data, biological sample size had less influence on TP identification, with the larger dataset identifying similar numbers of, or only slightly more, TPs than the smaller dataset (Figure 2a-e). The influence of sample size on the rate of identification of FPs differed between GEA programs. Larger sample sizes decreased the number of FPs identified by BayPass (and RDA for *Sim microcarpa*), but increased the number of FPs in LFMM2 analyses. This impacted the proportion of TPs in all identified associations for different sample sizes. For both simulated datasets, the proportion of TPs was higher in the BayPass analyses that had the larger sample sizes (higher TP:AA ratios), reflecting the opposing increases and decreases in TPs and FPs respectively. In LFMM2 analyses, as both TPs and FPs increased, sample size did not have a clear effect on the proportion of TPs.

### References

Bivand, R., Altman, M., Anselin, L., Assunção, R., Berke, O., Bernat, A., & Blanchet, G. (2019). spdep: Spatial dependence: Weighting schemes, statistics and models. R package version 1.1-2.

Griffith, D. A. (2005). Effective geographic sample size in the presence of spatial autocorrelation. *Annals of the Association of American Geographers*, *95*(4), 740–760.

VanDerWal, J., Falconi, L., Januchowski, S., Shoo, L., & Storlie, C. (2014). SDMTools: Species Distribution Modelling Tools: Tools for processing data associated with species distribution modelling exercises. *R package version*, *1*, 1-221.

**Table S1.** Spatial autocorrelation and effective sample size calculations for all climate variables. ESS = effective sample size; Emic = Eucalyptus macrocarpa; Ccal = Corymbia calophylla.

| Variable | MoransI_Emic | ESS_Emic | MoransI_Ccal | ESS_Ccal |
|---|---|---|---|---|
| BIO1 | 0.2717803 | 14.9703325 | 0.286601 | 14.9928764 |
| BIO2 | 0.12310717 | 20.9052364 | 0.18554264 | 18.9335779 |
| BIO3 | 0.32545538 | 13.165605 | 0.22685592 | 17.2414254 |
| BIO4 | 0.36919363 | 11.8189352 | 0.26051704 | 15.9460984 |
| **BIO5** | **0.2674845** | **15.1223146** | **0.32682604** | **13.6066236** |
| BIO6 | 0.09395951 | 22.222028 | 0.13972078 | 20.9413016 |
| BIO7 | 0.29576109 | 14.1426722 | 0.23091276 | 17.0813655 |
| BIO8 | 0.38415879 | 11.382467 | 0.14242746 | 20.8190133 |
| BIO9 | 0.32551934 | 13.1635562 | 0.32260337 | 13.7474863 |
| BIO10 | 0.3251226 | 13.1762678 | 0.36130331 | 12.4961223 |
| BIO11 | 0.13096385 | 20.5576531 | 0.17223335 | 19.5027759 |
| BIO12 | 0.16099944 | 19.2617443 | 0.14412258 | 20.7426558 |
| BIO13 | 0.13804338 | 20.2474132 | 0.12129118 | 21.7854966 |
| **BIO14** | **0.24401222** | **15.9730246** | **0.3149436** | **14.0057641** |
| BIO15 | 0.26701657 | 15.1389385 | 0.23603161 | 16.8809555 |
| BIO16 | 0.0820648 | 22.7695253 | 0.12771945 | 21.4888061 |
| BIO17 | 0.32200483 | 13.2764805 | 0.24931802 | 16.3688168 |
| BIO18 | 0.4330913 | 10.0373069 | 0.25460338 | 16.1682988 |
| BIO19 | 0.06752446 | 23.4435812 | 0.10887986 | 22.3646785 |

**Table S2**. Correlation between significant values (calibrated p-values for LFMM2 and Bayes Factor for BayPass) among the most conservative and liberal simulated datasets. Below the diagonal is correlation among LFMM2 significant values and above the diagonal is correlation among BayPass significant values. All values are significantly correlated at $p < 0.001$. MD = missing data; MAF = minor allele frequency.

<u>*Sim calophylla*</u>

| | **MD** | **10%** | | | **50%** | | |
|---|---|---|---|---|---|---|---|
| **MD** | **MAF** | **0.1** | **0.05** | **0.01** | **0.1** | **0.05** | **0.01** |
| **10%** | **0.1** | 1 | 0.824 | 0.821 | 0.838 | 0.841 | 0.830 |
| | **0.05** | 1 | 1 | 0.848 | 0.855 | 0.864 | 0.856 |
| | **0.01** | 0.998 | 0.999 | 1 | 0.852 | 0.857 | 0.853 |
| **50%** | **0.1** | 0.995 | 0.996 | 0.997 | 1 | 0.888 | 0.878 |
| | **0.05** | 0.994 | 0.996 | 0.997 | 0.999 | 1 | 0.880 |
| | **0.01** | 0.988 | 0.991 | 0.994 | 0.996 | 0.998 | 1 |

<u>*Sim microcarpa*</u>

| | **MD** | **10%** | | | **50%** | | |
|---|---|---|---|---|---|---|---|
| **MD** | **MAF** | **0.1** | **0.05** | **0.01** | **0.1** | **0.05** | **0.01** |
| **10%** | **0.1** | 1 | 0.840 | 0.841 | 0.843 | 0.837 | 0.845 |
| | **0.05** | 1 | 1 | 0.866 | 0.843 | 0.857 | 0.864 |
| | **0.01** | 0.999 | 1 | 1 | 0.845 | 0.863 | 0.867 |
| **50%** | **0.1** | 0.992 | 0.992 | 0.993 | 1 | 0.850 | 0.853 |
| | **0.05** | 0.991 | 0.992 | 0.993 | 1 | 1 | 0.862 |
| | **0.01** | 0.987 | 0.988 | 0.990 | 0.998 | 0.999 | 1 |

**Table S3.** The mean and standard deviation of significance between the MAF 0.01 and 0.1 within each data set for *Sim calophylla*. "Δ sig TPs" indicates how many True Positive SNPs become significant using a lower minor allele frequency (MAF), which were previously called not significant with the higher MAF. BF = Bayes Factor.

| | | BayPass | Δ sig TPs ^ | | LFMM2 | Δ sig TPs ^ | |
|---|---|---|---|---|---|---|---|
| Sample size | Missing data (%) | BF difference (mean ± SD) | 0.05 - 0.1 | 0.01 - 0.05 | log($P$ value) difference (mean ± SD) | 0.05 - 0.1 | 0.01 - 0.05 |
| 162 | 10 | 3.26 ± 2.54 | 0 | 0 | -2.78 ± 2.18 | -1 | 5 |
| 162 | 30 | 3.42 ± 2.97 | 1 | 4 | -2.93 ± 2.32 | 4 | 9 |
| 162 | 50 | 3.26 ± 3.08 | 6 | 1 | -2.58 ± 2.13 | 6 | 6 |
| 270 | 10 | 2.04 ± 2.40 | 1 | 1 | -4.04 ± 3.91 | 0 | 1 |
| 270 | 30 | 1.91 ± 2.65 | 0 | 0 | -3.74 ± 3.65 | 3 | 13 |
| 270 | 50 | 1.85 ± 2.49 | 0 | 0 | -3.63 ± 3.55 | 5 | 11 |

^ excludes TPs no longer in the dataset due to filtering using a greater MAF

**Figure S1**. Distribution of missing data in simulated datasets.

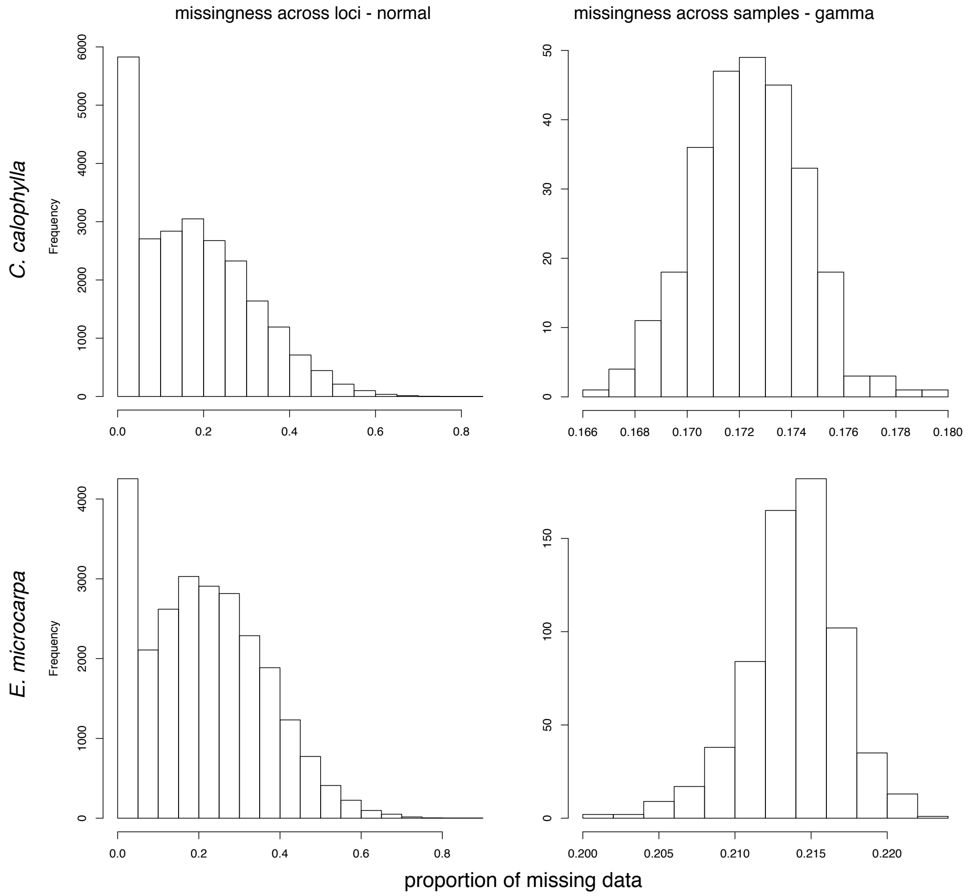

**Figure S2.** The relationship between total number of SNPs and the number of associations for *Eucalyptus macrocarpa* and *Corymbia calophylla*.

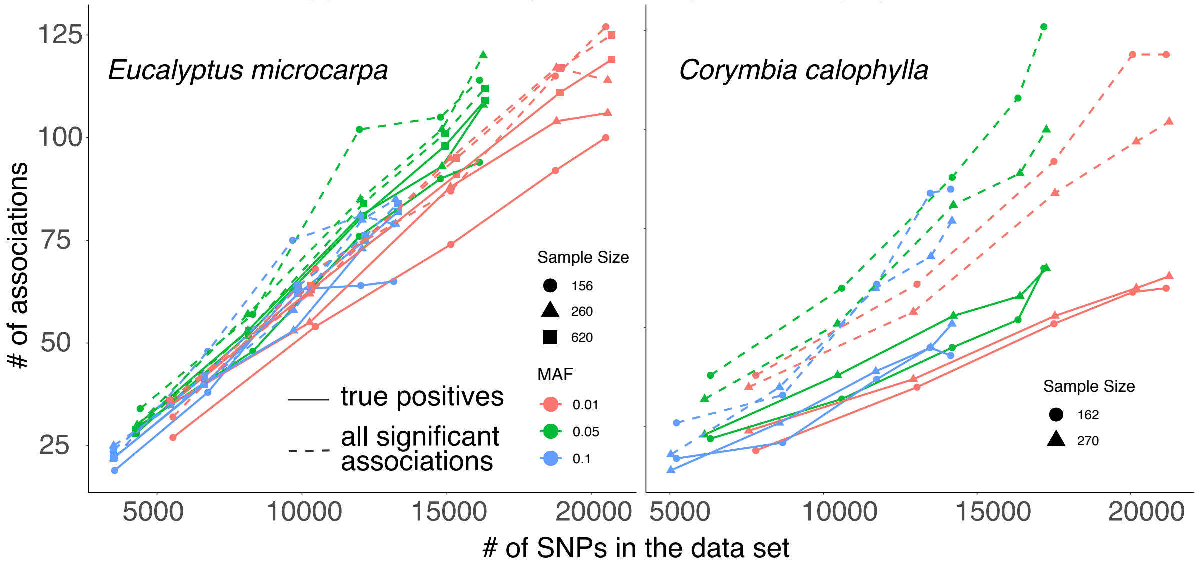

**Figure S3.** The relationship between strength of selection and significance for LFMM2 (Calibrated P-value) and BayPass (Bayes Factor).

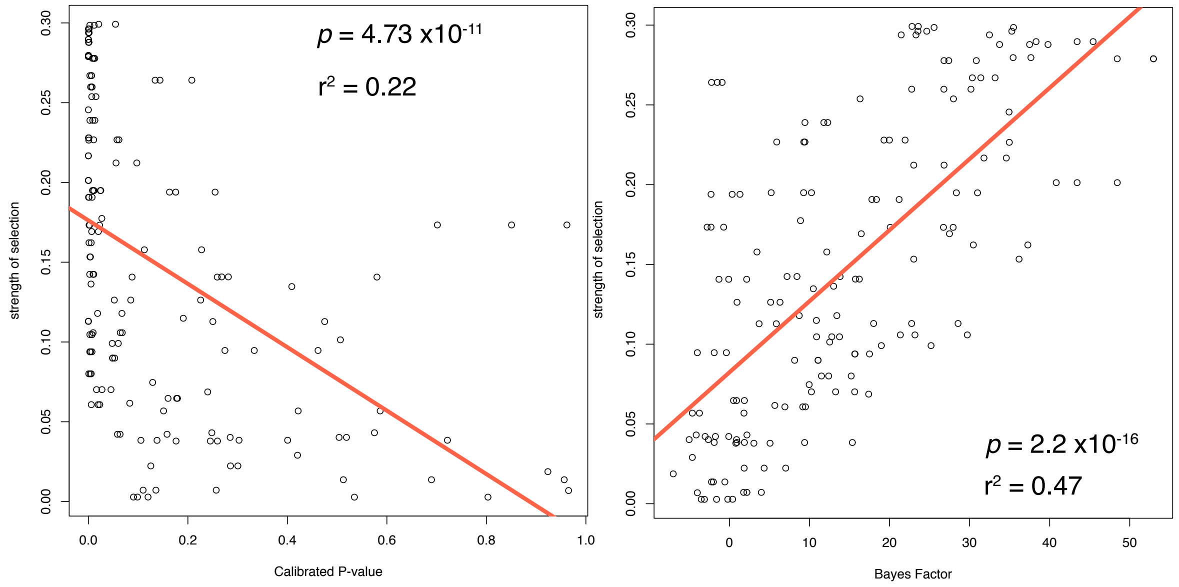

**Figure S4.** Effect of minor allele frequency (MAF), missing data (MD), and sample size (SS) on adaptive predictions. The difference between the two predictions are the most 'liberal' minus the most 'conservative' datasets. For MAF, MD is held constant at 30% and SS was held constant at 10 individuals per population (260 for *Sim calophylla*; 270 for *Sim microcarpa*). For MD, MAF was held constant at 0.05 and SS was held constant as above. For SS, MAF was held constant at 0.05 and MD was held constant at 30%.

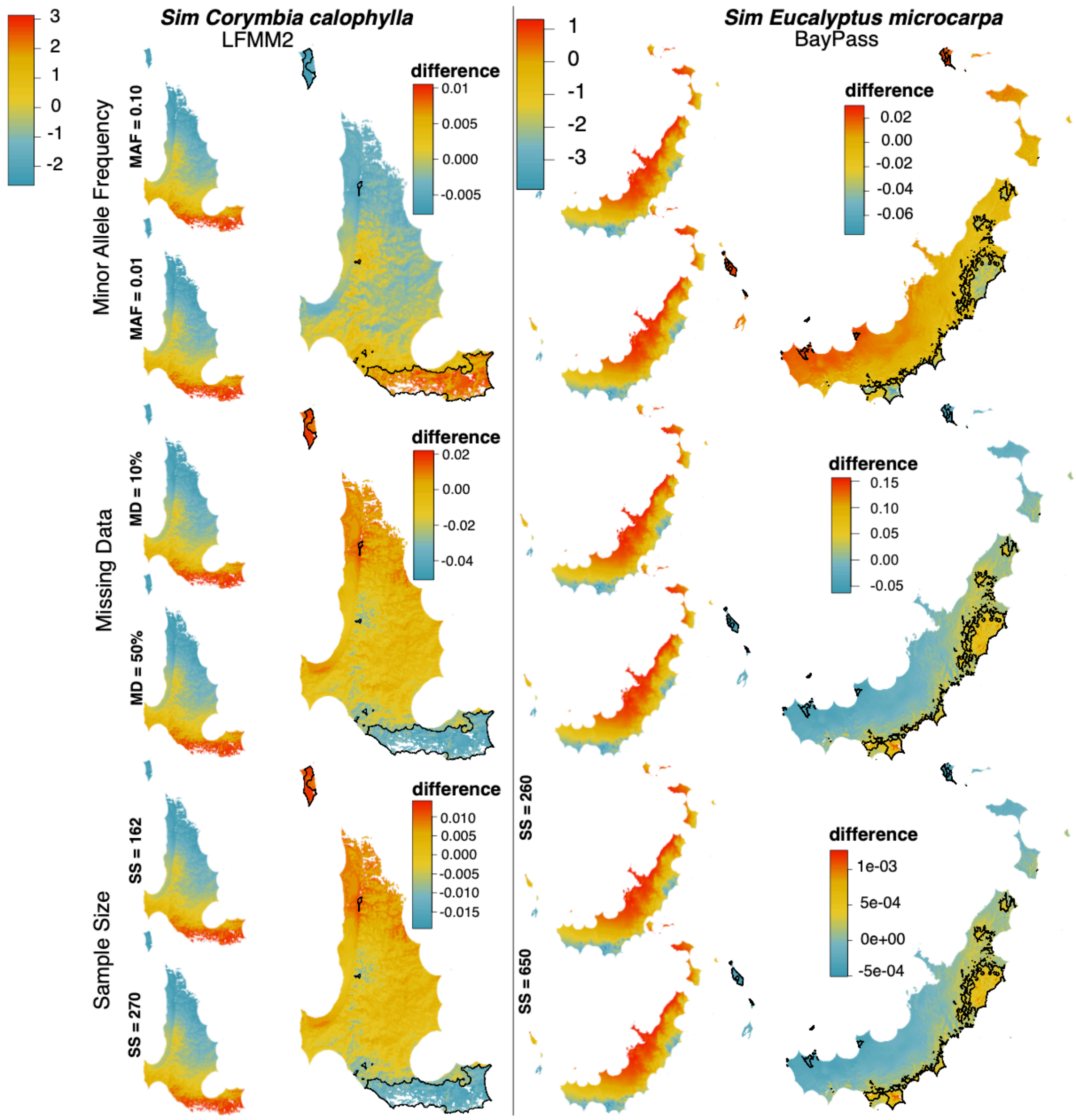
